# 3D bioprinting of Liver Microenvironment Model Using Photocrosslinkable Decellularized Extracellular Matrix based Hydrogel

**DOI:** 10.1101/2024.04.17.589849

**Authors:** Nima Tabatabaei Rezaei, Hitendra Kumar, Hongqun Liu, Ashna Rajeev, Giovanniantonio Natale, Samuel S. Lee, Simon S. Park, Keekyoung Kim

## Abstract

The liver, as one of the vital organs in the body, plays a crucial role in various bodily functions. Numerous factors can cause liver damage, that the sole remedy for severe liver conditions is the transplantation of healthy liver tissue. In response to the transplantation challenges, innovative approaches involving hydrogel-based technologies have emerged, leading to the creation of highly functionalized tissues. The development of three-dimensional printing and patterning of cell-laden biomaterial matrices offers promising advances for creating tissue-specific structures in tissue engineering and bioprinting. However, the matrix materials currently employed in bioprinting liver microtissue often fail to capture the complexity of the natural extracellular matrix (ECM), hindering their ability to restore innate cellular shapes and functions. Liver ECM-based hydrogels are increasingly recognized for their potential as biomimetic 3D cell culture systems that facilitate the exploration of liver disease, metabolism, and toxicity mechanisms. Yet, the conventional production of these hydrogels relies on slow thermal gelation processes, which restrict the manipulation of their mechanical characteristics. In this research, we introduce a novel approach with a functionalized photocrosslinkable liver decellularized extracellular matrix (dECM). By combining liver dECM methacrylate (LdECMMA) with gelatin methacrylate (GelMA), we achieved accelerated crosslinking under visible light irradiation and the ability to tune the mechanical, rheological, and physiological properties of the material. We encapsulated human hepatocellular carcinoma cells within an optimal concentration of the GelMA-LdECMMA hybrid hydrogel and examined cell proliferation and function over an extended period. The findings revealed that the GelMA-LdECMMA hybrid hydrogel enhances liver cell proliferation and function, holding significant promise for applications in drug screening and liver cancer metastasis research.

## 1. Introduction

The liver is an essential organ responsible for maintaining the body’s biochemical balance, including the synthesis of blood proteins, glucose metabolism, and the detoxification of various substances [1–3]. Despite its critical functions, the incidence of liver disease-related fatalities is on the rise globally [4]. Liver transplantation or cell therapy techniques such as hepatocyte transplantation [5] has shown promising potential to be used as viable treatment for end-stage liver disease but is limited by the scarcity of donor organs and challenges with cell engraftment [6]. Consequently, addressing end-stage liver disease remains a significant hurdle, filled with uncertainties regarding effective treatment methods. In recent times, advancements in bioengineering, particularly through tissue engineering techniques, have led to the exploration of new solutions. These include the creation of transplantable liver microtissues or entirely bioengineered livers, offering promising alternatives for the long-term restoration of liver function [2,7].

As the newly emerged class of biomaterials, tissue engineering approaches have effectively harnessed the decellularized extracellular matrix (dECM) to mimic the native cellular microenvironment for tissue fabrication *in vitro* [8,9]. The ECM’s unique three-dimensional (3D) microstructure and organization are pivotal in regulating cell behavior, communication, and maintaining tissue integrity and functionality [10]. A prevalent method for obtaining dECM employs various chemical agents, including ionic and non-ionic detergents, acids and bases, alcohols, and chelating agents [11]. By integrating these tissue-specific extracellular matrix (ECM) components into 3D model systems, it is possible to facilitate the development of tissues that more accurately resemble their physiological counterparts in terms of function and structure [12].

Structure of the fabricated scaffolds, along with the composition of the scaffold biomaterial, plays an equally important role in defining the microenvironment and, consequently, the function of the fabricated tissues. While progress has been made in liver tissue engineering using extrusion-based bioprinting techniques, these efforts are hindered by challenges such as low printing resolution and compromised structural integrity [13]. These issues stem from the inherent limitations of extrusion-based printing methods and the inadequate mechanical properties of decellularized extracellular matrix (dECM)-based bioinks [14]. To address these limitations, photocuring-based biofabrication methods, including digital light processing (DLP) [15,16] have been employed to create liver models for various biomedical applications. DLP printing can achieve high-resolution hydrogel structures with intricate architectures and can enhance the mechanical properties of the printed hybrid and composite hydrogels [16]. However, DLP printing involves the use of potentially harmful photoinitiators that, along with UV light, can unpredictably affect cell behavior. As a result, recently, there has been a growing interest in exploring photoinitiators that activate under visible light excitation [17].

In order to swiftly produce hydrogels that exhibit controlled gelation processes, the technique of functionalizing natural polymers with photocurable components has been utilized [18–20]. This method combines the inherent benefits of natural polymers—such as biocompatibility and biodegradability—with the consistent physicochemical properties provided through chemical functionalization. The resulting hydrogels are capable of forming controllable, reproducible, and biologically relevant three-dimensional tissue models. However, existing ECM-based hydrogels for liver tissue have limited ability to manipulate their physical characteristics, and there have been relatively few attempts to enhance the physicochemical properties of liver dECM hydrogels [21,22].

Addressing the critical need for an enhanced liver matrix-derived bioink that guarantees superior printability, cell viability, and functionality, this study adopts a modified decellularization protocol for porcine liver using widely recognized chemical detergents like sodium dodecyl sulfate (SDS) and Triton X-100. The decellularized liver tissues were then solubilized and methacrylated, resulting in the formation of liver dECM methacryloyl (LdECMMA). This was further blended with gelatin methacryloyl (GelMA) to create hybrid hydrogels. upon photocrosslinking. By integrating LdECMMA with GelMA, we were able to bioprint liver microtissues with adjustable and enhanced physical and mechanical attributes. The LdECM-based hydrogels underwent comprehensive characterization to assess their functionalization degree, physical characteristics, and cytocompatibility. Ultimately, by encapsulating liver cells within the developed scaffold and monitoring their functionality over extended culture periods, we demonstrated the scaffold’s efficacy towards liver tissue engineering applications.

## 2. Materials and Methods

### 2.1 Porcine Liver Decellularization

Fresh porcine liver tissues were purchased from a local butcher shop and transported on ice to the lab and stored in −80 °C freezer prior to decellularization process. Previously reported protocols were modified to prepare the liver decellularized extracellular matrix (dECM) [23–25]. In brief, frozen liver tissues were thawed and cut into small cubic pieces (less than 1mm) with scissors and washed in distilled water for 3 h to remove blood. Next, the samples were soaking in 1% (w/v) Triton X-100 (VWR, Mississauga, ON, Canada) for 24 h, followed by treatment with 0.1% (wt/vol) sodium dodecyl sulfate (SDS) (VWR, Mississauga, ON, Canada) solution for 48 h, replacing fresh solution after first 24 h. A solution to liver tissue ratio of 10:1 was maintained through all the washing steps. Then, tissue pieces were rinsed thoroughly with distilled water several times and blended with high speed to have uniform product. Finally, blended liver tissue was separated by centrifuging at 3500 rpm for 3 min and was stored at −80 °C and then lyophilized to obtain final dECM.

### 2.2 Decellularization Assessment

To visualize and investigate residual cells and microarchitecture, both native and decellularized porcine liver tissues were fixed in 4% (wt/vol) paraformaldehyde solution at room temperature for 24 h followed by sequential immersing in 100%, 95%, 90%, 75%, 70% and 50% (vol/vol) ethanol overnight, and then embedded in paraffin.

Paraffined sections were cut into thin slices of 5-7 μm thickness for further staining steps. Then, tissue sections were stained by hematoxylin and eosin (H&E) (VWR, Mississauga, ON, Canada) and Masson’s trichrome staining (Polysciences Inc., Warrington, PA, USA) in order to observe cell component residues and collagen distribution, respectively. The H&E and Masson trichrome staining samples were examined using an inverted microscope (ECHO revolve).

Slices from the liver specimens and the dECM samples were subjected to DAPI staining. This technique was employed to visualize the nuclei within the liver tissues, and to assess the presence or absence of nuclei in the dECM samples, respectively. Fluorescence microscopy images were acquired using an inverted fluorescence microscope (ECHO revolve) using the DAPI fluorescence channel (*λ_ex_350nm/λ_em_465nm*).

DNA and total protein concentration quantified content were measured using the PicoGreen dsDNA Assay Kit (ThermoFisher, Waltham, Massachusetts, USA) and TPC kit (ThermoFisher, Waltham, Massachusetts, USA), respectively, to indicate residual cells and protein content in the dECM (n=3). Briefly, 50 mg of native liver and lyophilized liver dECM were added to 1 ml of pepsin digest solution (Ward’s Science, ON, Canada) with a concentration of 0.1 mg/ml and homogenized with homogenizer (FSH-2A High Speed Homogenizer) at 20000 rpm. Then samples were vortexed and kept in 65 °C water bath for 6 h followed by centrifuging at 4000 rpm for 15 min to collect the supernatant. A microplate reader (SpectraMax ® iD3 Microplate Readers) was used for fluorescence intensity measurement 480 nm excitation and absorbance measurement under 560 nm for DNA and TPC kits, respectively.

### 2.3 dECM functionalization

Lyophilized dECM was digested in 0.5 M of acetic acid (MilliporeSigma, Oakville, ON, Canada) and 10 mg pepsin solution per 100 mg dECM for 3 days at room temperature in order to obtain uniform solution. After complete digestion, using 1 M NaOH (MilliporeSigma, Oakville, ON, Canada) solution, the dECM solution’s pH was adjusted approximately 8.5. Then, 300 µl of glycidyl methacrylate (MilliporeSigma, Oakville, ON, Canada) per 0.5 g dECM in the solution was added dropwise at room temperature and agitated aggressively for 24 h. The resulting biopolymer (dECMMA) was dialyzed in a 12–14 kDa dialysis tubing (ThermoFisher, Waltham, Massachusetts, USA) against distilled water for 3 days with water changed twice a day. Then, the solution was lyophilized at −84 °C for another 3 days. The lyophilized dECMMA were stored at −80 °C for further use.

The degree of substitution of the synthesized LdECMMA was measured using proton nuclear magnetic resonance (1H NMR) as reported previously [26,27]. LdECMMA solution was prepared with a concentration of 0.5% (w/v) 1 mL of deuterium oxide (ThermoFisher, Waltham, Massachusetts, USA). Next, a 600 MHz NMR spectrometer (Bruker?) was used to record the 1H NMR spectra for the synthesized LdECMMA samples. The internal reference was set to hydroxyl signals (0.5-1 ppm). The peaks of primary amine groups (2.8–2.95 ppm) were integrated to compute the degree of substitution.

### 2.4 Preparation of prepolymer solution and bioprinting system

GelMA was synthesized following a previously reported [28]. In brief, 5 g of powdered gelatin from porcine skin (Type A, Bloom strength 300, Sigma-Aldrich, St. Louis, MO, USA) was dissolved in 50 mL of RO purified water. After complete dissolution at 48-50 °C, 9-10 ml of GMA (MilliporeSigma, Oakville, ON, Canada) was added to the gelatin solution dropwise. The solution was then maintained at 48-50 °C with constant stirring at 750 rpm for 12 h. Upon completion of the reaction, the solution was transferred into dialysis tubing (12–14 kDa) to be dialyzed against distilled water for 3 days with water changed twice a day. After 3 days, the remaining solution was frozen and then lyophilized to obtain a foamy solid of GelMA. The lyophilized samples were stored at −20 °C for further use. In order to prepare hybrid prepolymer solution, 5% (w/v) GelMA and different concentrations of LdECM or LdECMMA were dissolved in PBS, as listed in **Table 1**. For this purpose, first 5% (w/v) GelMA solution was prepared and then different amounts of LdECM or LdECMMA was added to the solution and stirred overnight at room temperature to achieve a homogeneous prepolymer solution. Next, the photoinitiators for visible light crosslinking were added to the prepolymer solutions. An eosin Y (EY)-based two-component photoinitiator as reported in earlier studies was used for the crosslinking of prepared bioinks [29,30]. The two-component photoinitiator was made up of 0.02 mM EY disodium salt (VWR, Mississauga, ON, Canada) and 0.2% w/v triethanolamine (TEOA) (MilliporeSigma, Oakville, ON, Canada). After preparing LdECMMA-GelMA solution, EY and TEOA were added to reach the final concentration of 1% (v/v) for each component. The designed structures were formed by exposing the prepolymer solutions to visible light (wavelength: 400–700 nm, intensity: 80.8 mW.cm^−2^) in the modified custom made DLP bioprinting system reported earlier [29,31]. The exposure duration was varied for each bioink combination based on their photocrosslinking characteristics. Circular vats with 500 µm depth were used to crosslink the bioinks and form the designed structures in one step.

**Table 1.**
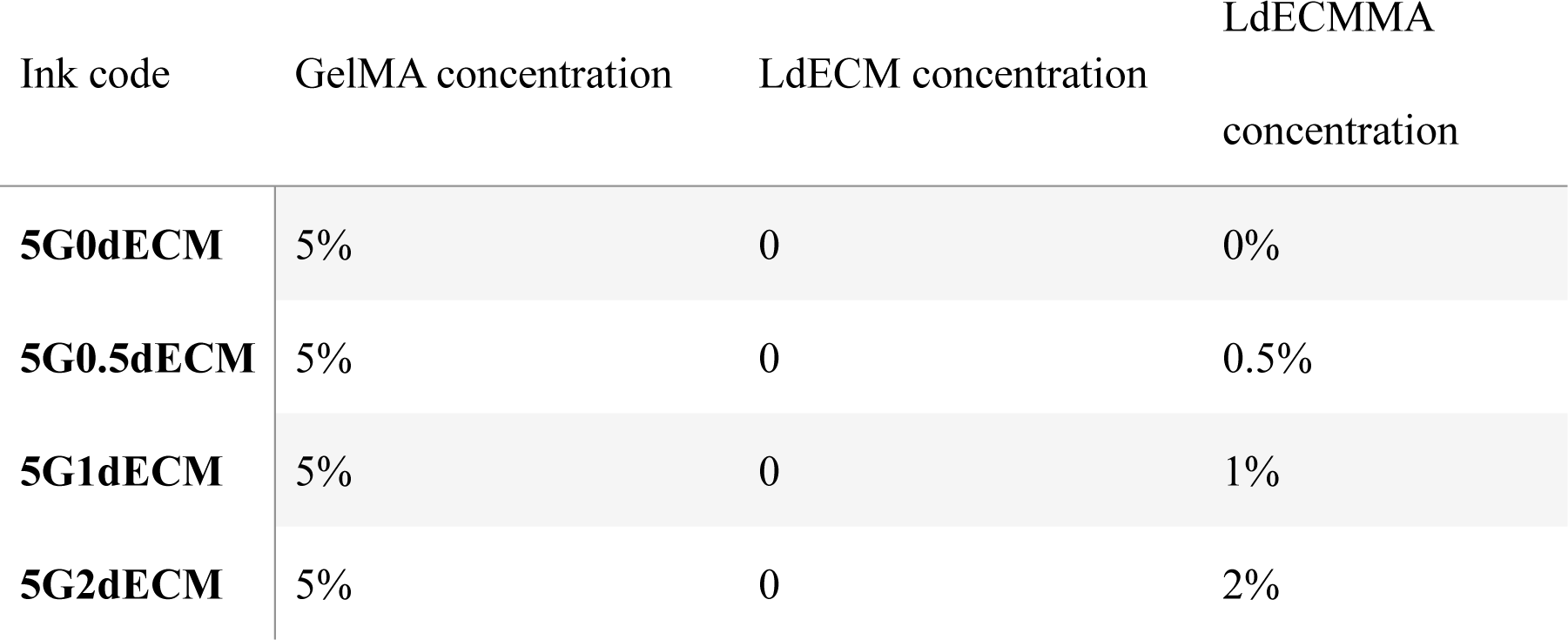
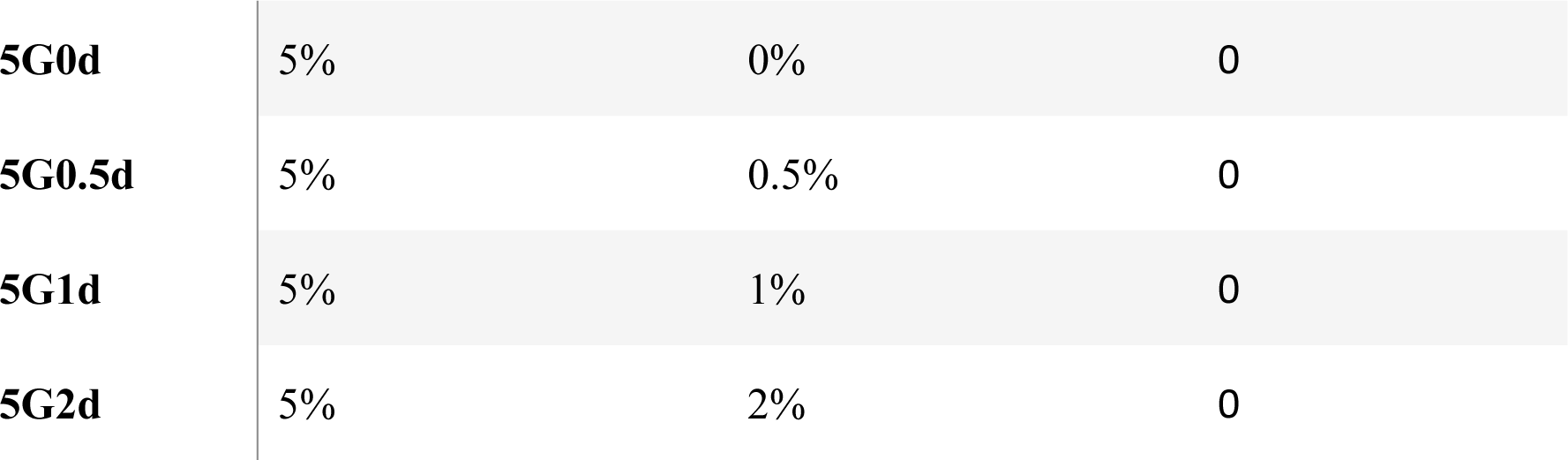
Different compositions of dECM-based bioinks.

### 2.5 Measurements of mechanical, swelling and degradation properties

The mechanical properties of the hydrogels were assessed by evaluating the compressive modulus of the crosslinked hydrogels as a measure of the mechanical stiffness. The probe used for the compression was a custom-made flat-ended rigid cylinder with 12.5 mm diameter. In order to prepare samples, a 2 ml solution of the hydrogel precursor with different concentration of LdECMMA was prepared and transferred into cylindrical molds with 8 mm diameter and 8 mm depth. A prepolymer solution volume of was added in each well. The prepolymer solution was then crosslinked in the SLA bioprinting system for 5 min due to the large volume using the SLA bioprinting system to ensure the whole volume was crosslinked. Additionally, to observe the influence of functionalization of dECM on the mechanical properties of the hybrid hydrogels, unmodified LdECM was also used to prepare prepolymer solutions with the same concentration and crosslinked under same conditions as GelMA-LdECMMA prepolymer solutions. Afterwards, a vertical axis micromechanical testing machine along with the custom flat-cylindrical probe was used to record the force vs. displacement plots for hydrogels under compression. Up to 70% compression was applied on the hydrogel surface to obtain the force-displacement plot. Using initial dimensions of the hydrogel samples and a custom-made MATLAB script, compression modulus was determined using initial 10% of strain area slope (linear region).

The swelling ratio of the crosslinked hydrogel samples was measured to evaluate the water uptake capability. To describe the procedure briefly, 250 µL of different conentrations of hydrogel solutions were casted in 8 mm diameter and 8 mm deep cylindrical mold and crosslinked under visible light using the DLP 3D bioprinter for 5 min. Subsequently, the crosslinked hydrogels were immersed in PBS and kept in an incubator at 37 °C and 5% CO2for 24 h. Each sample’s hydrated weight (ww) was recorded at hydration equilibrium. Afterwards, they were frozen in a −80 °C freezer and lyophilized for 48 h before their dry weight (wd) was measured. Finally, the following equation was used to calculate the swelling ratio (n=5):

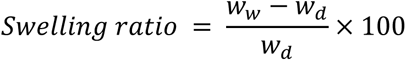

In order to investigate the hybrid hydrogels’ degradation behaviour, 750μl of LdECMMA-GelMA hydrogels were cast in 96-well plates and cured for 5 mins under visible light. In addition, a sample containing 5% GelMA and 0.5% unmodified and solublized LdECM was used to demonstrate the influence of methacrylation on the degradation properties of the hydrogels. All the samples were lyophilized for 48 h and weighed to get the initial weight (w0), and then were immersed in PBS at 37 °C for 24 h.

The swollen samples were placed in a 48-well plate with collagenase-I enzyme solution (50 U/mL) and incubated at 37 °C for hydrogel degradation study. Three samples were collected every 1 h, frozen at −80°C and lyophilized for 24 h. The lyophilized samples were weighed to obtain the remaining polymer matrix weight (wr) after degradation. The biodegradation property (Qd) of hydrogels was indicated by the percentage of the remaining weight of the disks after degradation as:

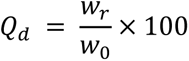

### 2.6 Scanning Electron Microscopy (SEM) and porosity quantification

Scanning electron microscopy (SEM) was used to investigate the microscale structure of the LdECMMA-GelMA hybrid hydrogels,. For this characterization, 250 µL prepolymer solution was prepared for each hydrogel group at defined concentrations and, crosslinked with visible light irradiation. Subsequently, the samples were frozen in a −80 °C freezer followed by lyophilization for two days. The lyophilized samples were then sputter-coated with carbon sputtering machine and imaged using SEM (Phenom Pro X).

### 2.7 Crosslinking kinetic evaluation and Rheological Characterization

The rheology characteristics of GelMA and different types LdECMMA-based hybrid bioinks were evaluated using a rheometer. All measurements were performed at room temperature using a 25 mm diameter parallel plate geometry and maintaining a 0.5 mm gap. The viscosity variation of the bioinks was recorded by varying the shear rates from 0.01 to 100 s^−1^. The viscosity characterization of the bioinks was performed in absence of visible light irradiation. For the photocrosslinking kinetics evaluation, the bottom plate is replaced with a transparent surface to allow light irradiation during the crosslinking process. When recording the photocrosslinking kinetics, the visible light was irradiated to the sample from the bottom to initiate the crosslinking as shown in **Figure 3(B)**. A visible light was projected 1 min after the commencement of the experiment and the exposure continued for a duration of 5 min. The evolution of the rheological properties of the gel during photocuring was tracked in oscillatory mode at constant shear rate of 1% and angular frequency of 1 Hz. The storage and loss modulus for the photocrosslinking bioink were recorded simultaneously. After completing the exposure duration, angular frequency sweep was done in the range of 0.1-100 rad/s at constant 10% strain to observe dynamic rheological properties.

**Figure 1.**
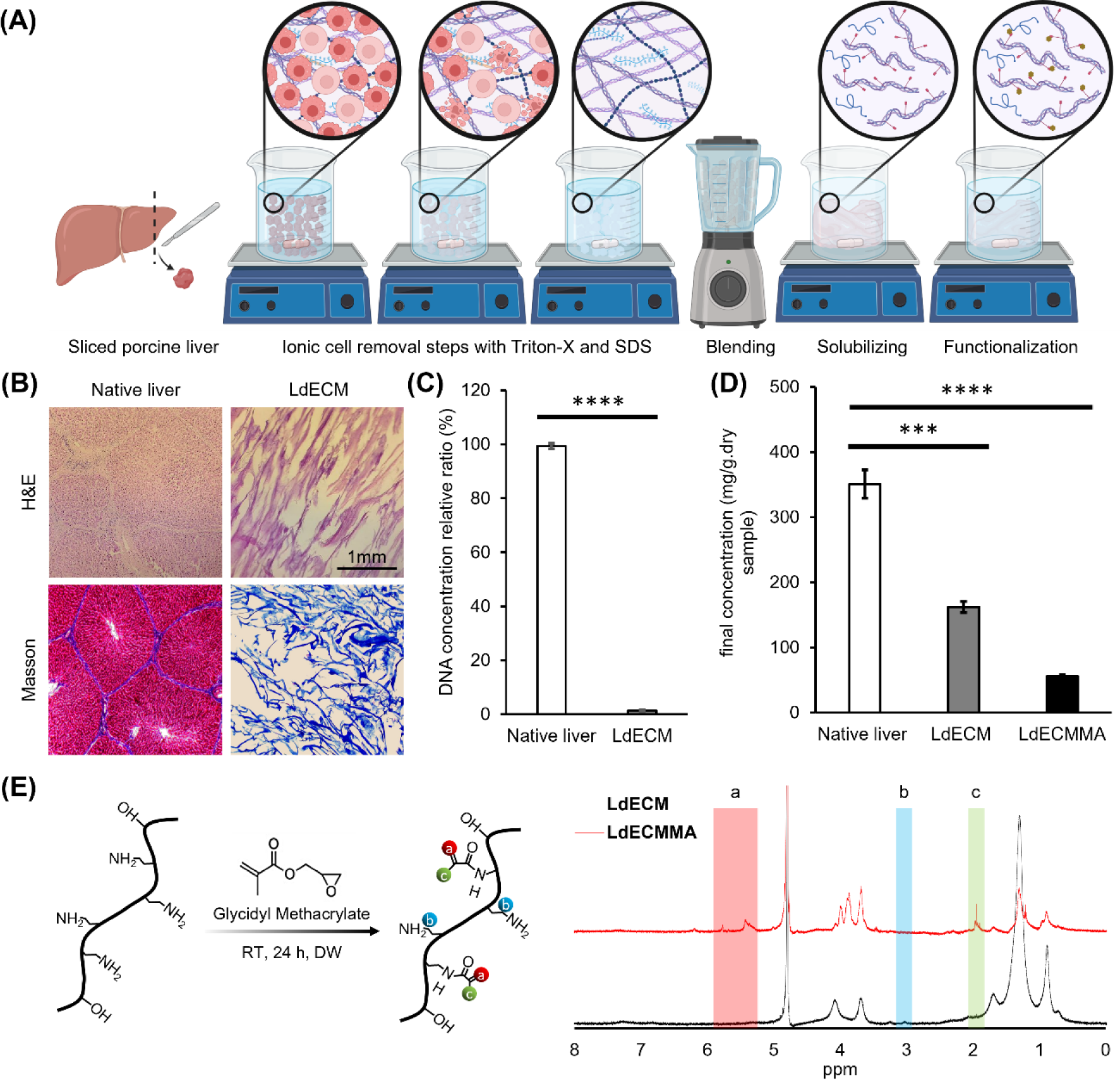
Liver dECM synthesis and methacrylation (LdECMMA) process and relevant characteriztions. (A) Schematic representing the washing steps, solubilizing, and functionalization steps. (B) H&E and Masson Trichrome staining of native liver tissue and decellularized liver tissue ECM. Biochemical characterization before and after decellularization (C) total DNA content evaluation. (D) Total protein concentration of native liver, liver dECM and liver dECMMA. (E) The reaction mechanism of substitution of methacrylate groups on dECM backbone molecules, and degree of methacrylation characterization of LdECMMA using ^1^H NMR spectra of methacrylated and unmodified LdECM. (* p < 0.05, ** p < 0.01, *** p < 0.001 and **** p < 0.0001)

**Figure 2.**
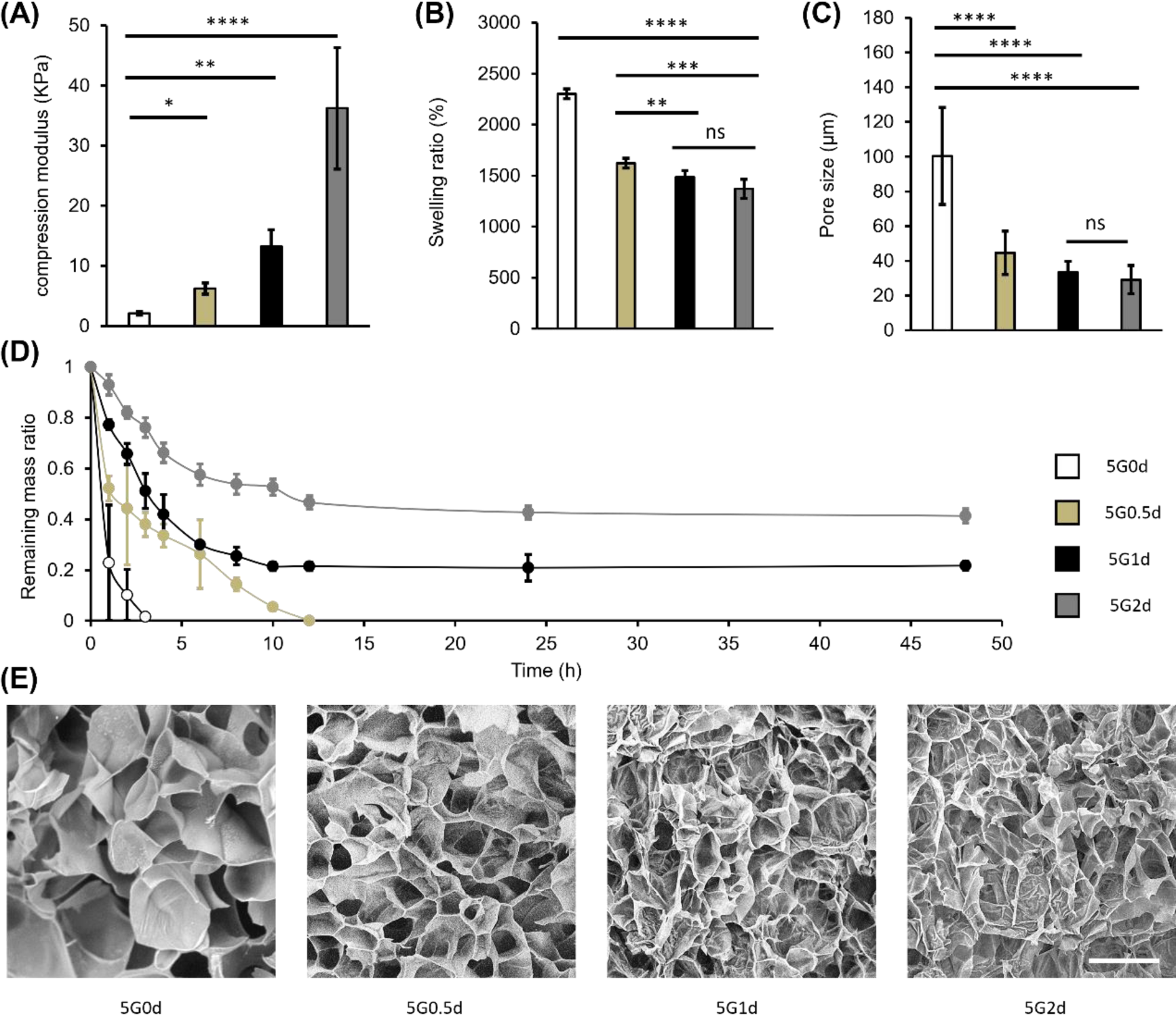
Mechanical and physical characterization of hybrid GelMA-LdECMMA hydrogels. (A) Compression modulus different compositions of hydrogels (n=5). (B) Mass swelling ratio of different compositions of hydrogels (n=5). (C) Pore size evaluation of the 3D microstructure of the hydrogels (n=5). (D) Degradation characterization of the hydrogels under enzymatic degradation (n=5), (E) Scanning Electron Microscopy images showing porous structure (scale 100 µm) (* p < 0.05, ** p < 0.01, *** p < 0.001 and **** p < 0.0001)

**Figure 3.**
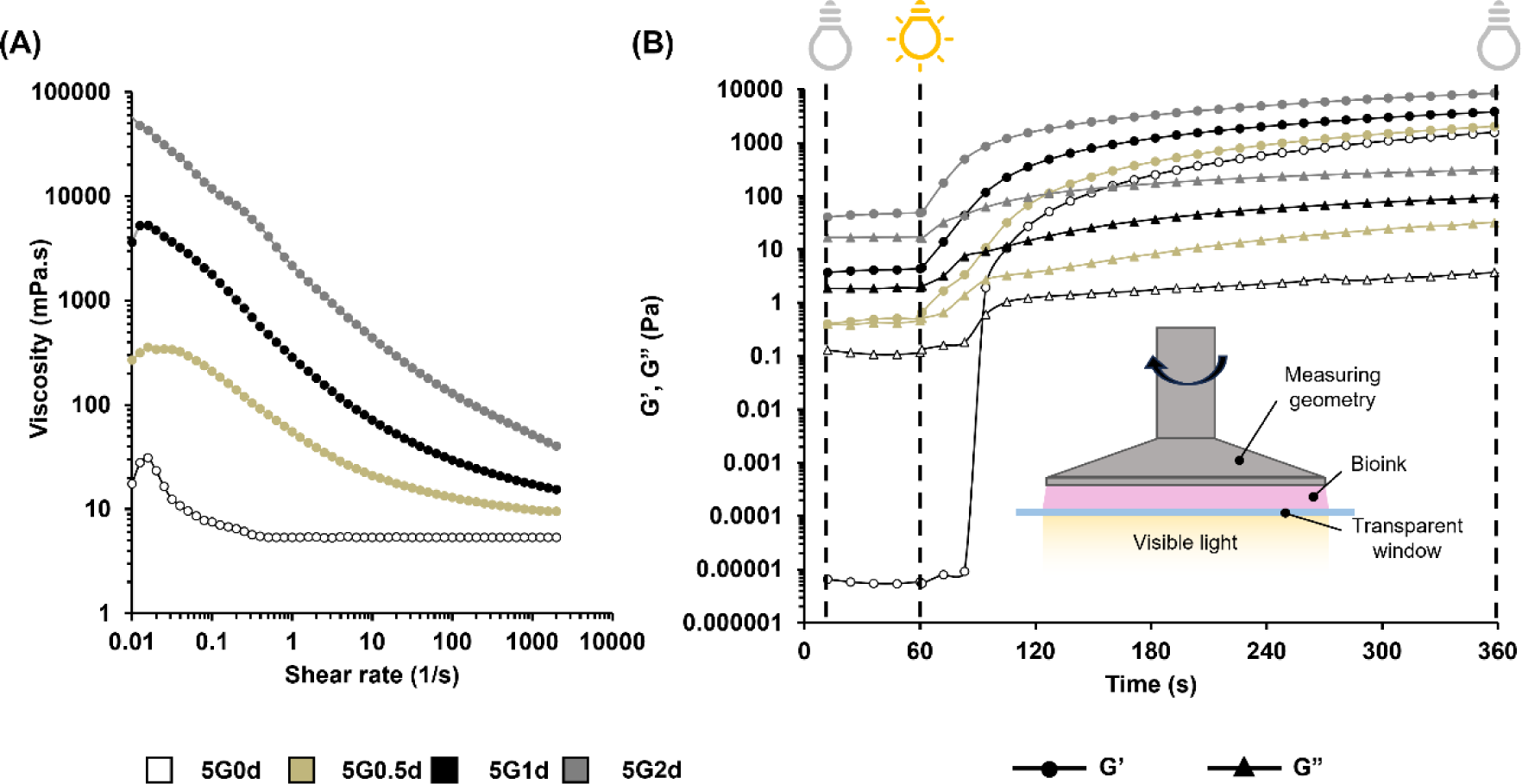
Dynamics rheological characterizations of GelMA and LdECMMA containing hydrogels. (A) Comparison of viscosity vs. shear rate for different compositions of dECM-based hydrogels. Photocuring kinetics of GelMA and GelMA-LdECMMA hydrogels represented by (B) storage modulus (G’) and loss modulus (G”) recorded in situ crosslinking, for all in situ crosslinking characterizations, light irradiation started at t=60 s and ended at t=360 s.

### 2.8 Cell culture

Human hepatocellular carcinoma HepG2 cells were cultured utilizing high glucose Dulbecco’s Modified Eagle Medium (DMEM) (Corning, Arizona, USA) supplemented with 10% fetal bovine serum (FBS) (Corning, Arizona, USA) and 1% penicillin/streptomycin (Cytiva, Marlborough, MA, USA). For culturing the HepG2 cells, the cells were cultured in tissue culture flasks and maintained at 37 °C and 5% CO2 in an incubator. The growth medium was renewed every 3 days.

### 2.9 Cell-laden hydrogels viability assessment

Cell viability analysis was conducted utilizing the LIVE/DEAD Viability/Cytotoxicity Kit (Biotium, Fremont, CA, USA). For cell viability and growth assessments, the cells were seeded at a density of 2.6 × 10^6^ cells mL^-1^ in various bioinks and the crosslinked structures were cultured in the same growth medium renewing every 3 days.

Following 3 washes with PBS, the encapsulated cells were subjected to staining for 30 minutes in the dark at room temperature using a PBS solution containing 0.5 µL mL^−1^ calcein AM and 2 µL mL^−1^ EthD. Subsequently, the samples underwent 3 additional washes with PBS to eliminate any remaining reagents. Fluorescence images were captured using an inverted fluorescence microscope, with Calcein AM fluorescence recorded in the FITC channel and EthD fluorescence in the TxRed channel.

### 2.10 Metabolic activity assay

Cellular mitochondrial metabolic activity was assessed by adding the oxidation-reduction indicator, tetrazolium hydroxide salt assay (XTT kit; Biotium, Fremont, CA, USA), at a ratio of 1/10 of the medium volume to evaluate cell proliferation. Following a 24-hour exposure to XTT solution at 37°C in a 5% CO2 environment, 100 μl of medium from each well was transferred to a 96-well plate. The adsorption metabolite of resazurin, resorufin, was then measured using a spectral scanning plate reader with excitation at 450 nm.

### 2.11 Proliferation and morphological assay

To evaluate cell proliferation in hydrogel scaffolds containing either control or LdECMMA bioinks, cells were encapsulated and cultured for one month. A hydrogel prepolymer solution was prepared with 5% w/v GelMA (5G) (used as the control) and 5% w/v GelMA supplemented with 0.5% LdECMMA (5G0.5d). Cells were added at a density of 2.6 × 10^6^ cells ml^−1^ and evenly dispersed. The produced bioink served as the ink for the custom bioprinter described in an earlier report [32]. Multiple disks, each with a diameter of 4 mm and a thickness of 400 μm, were printed to ensure that at least one dimension fell in the range comparable to the critical limit for media diffusion. After crosslinking with visible light for 1 minute in a custom-made DLP bioprinting system [32], the disks were transferred to a petri dish, washed with PBS, and incubated in fresh growth media at 37°C with 5% CO2. To assess cell morphology and proliferation, at least three disks per sample were stained with phalloidin (Cytoskeleton Inc., Denver, CO, USA) and DAPI (MilliporeSigma, Oakville, ON, Canada) for cytoskeleton and nuclei, respectively, at 7, 14, 21, and 28 days of culture. Briefly, samples were fixed with paraformaldehyde, permeabilized with Triton X-100, stained with phalloidin 488, and DAPI, and imaged using an inverted fluorescence microscope equipped with DAPI and FITC channels.

### 2.12 Immunostaining and fluorescent microscopy

Cell encapsulation and sampling was done according to the procedure mentioned in section 2.10. For immunofluorescence staining of Albumin, the manufacturer’s protocol for antibodies was followed. Initially, 3D cultured HepG2 cells were rinsed with PBS to remove excess media absorbed by the hydrogels. Subsequently, the structures were fixed in 4% paraformaldehyde in PBS for 90 minutes at room temperature. Following three washes with PBS containing 0.3% Triton X-100 (washing solution), the cells were blocked with 1% Bovine Serum Albumin (BSA) (MilliporeSigma, Oakville, ON, Canada) and 0.3% Triton X-100 in PBS for 1 hour to minimize nonspecific binding. After rinsing with the washing solution, cells were incubated with primary antibodies: rabbit anti-albumin antibody (ab207327; Abcam, Toronto, ON, Canada) diluted 1/500 in PBS at 4°C overnight. Following incubation, 3D cultured cells were washed with the washing solution and then incubated with goat anti-rabbit secondary antibody Alexa Fluor 488 (ab150077; Abcam, Toronto, ON, Canada) for 2 hours at room temperature. After three washes with PBS, the samples were transferred on a glass slide and mounted with the mounting solution containing DAPI. A coverslip was placed on top, and the samples were imaged using an inverted fluorescence microscope (ECHO revolve) equipped with DAPI and FITC channels.

### 2.13 Albumin secretion and metabolic analysis

To evaluate the liver-specific functions of cells within the hydrogel, secreted albumin levels were quantified using an Albumin ELISA kit (AFG Scientific, Northbrook, IL, USA), following the provided manufacturer’s instructions. Media was changed 24 hours before sampling. Collected media samples were stored at −20°C until use and subsequently measured after appropriate dilution (1:5) to ensure alignment with the kit’s standard curve.

### 2.14 Statistical analysis

Quantitative data are reported as means ± standard deviations (SD), with inferential statistics (p-values) used for analysis. Statistical significance was determined using two-tailed t-tests and ANOVA, with significance levels set at p < 0.05, p < 0.01, or p < 0.001. Analysis was conducted using GraphPad software.

## 3. Results and Discussion

### 3.1 Decellularization Assessment

In order to develop dECM-based bioinks capable of being used for DLP bioprinting, Porcine fresh liver was used for liver decellularization and functionalization, combined with gelatin methacrylate (GelMA). Through the process cells were washed out undergoing sequential steps utilizing ionic and non-ionic detergents, pepsin solubilizing and functionalization, followed by freeze drying (**Figure 1(A)**, **Figure S1**). The process of removing the cells was optimized to preserve native liver ECM key components such as collagen strands as well as cell elimination. Histochemical staining was performed to visualize cell nuclear material and collagen content. As shown in **Figure 1(B)**, H&E stained section of native liver revealed intact liver tissue morphology showing the cells arrangement in hepatic lobules. In contrast, H&E staining of the dECM samples showed collapsed morphology of the lobules with no visible cell nuclei (stained in dark purple as shown in native liver tissue) proving successful cell removal. Additionally, Masson staining was performed to determine the presence of the collagen in obtained dECM. The result revealed that the main component remaining after decellularization steps is the collagen fibers as stained in blue. Furthermore, the functionalized dECM material (LdECMMA) underwent Masson’s trichrome staining to assess the preservation of collagen fibers post-functionalization. **Figure S2** illustrates the prevalence of collagen fibers within the functionalized dECM sample. Nonetheless, when compared to **Figure 1(B)**, it is evident that the collagen fibers have become fragmented as a consequence of the solubilization process. Moreover, DAPI staining of both liver and dECM samples, as presented in **Figure S3**, serves as additional evidence of successful cell removal.

To evaluate the decellularization efficacy quantitatively, we investigated the DNA and total protein content. It is observed that the DNA content in dECM sample was dropped to less than 2.7 ± 0.31% of the DNA content in native liver (normalized to native liver DNA content) as presented in **Figure 1(C)**. Although ionic treatments have high efficacy on cell removal, they are more aggressive than non-ionic and will wash away more contents of ECM proteins. For investigating the influence of the washing and functionalization step on total protein concentration. it has been shown that the total protein concentration has been decreased significantly from 350.96 ± 21.66 to 162.05 ± 8.58 mg/g of dry sample (**Figure 1(D)**) which was predictable due to using ionic harsh detergent (SDS) capable of promoting protein erosion [33].

### 3.2 dECM functionalization and characterization

Moreover, the widespread utilization of self-assembled ECM-based hydrogels was hindered by their slow and uncontrolled gelation process, as well as their mechanical properties that do not mimic physiological conditions [34]. Presently available ECM hydrogel-based liver tissue models lack sufficient control over their physical characteristics [35,36], failing to adequately consider the relationship between hydrogel stiffness and liver physiology and pathology [37]. To address these challenges, researchers have explored the functionalization of natural polymers using photocurable components as a means to rapidly produce hydrogels with controlled gelation properties [38–40]. By combining the benefits of natural polymers, such as biocompatibility and degradability, with the reproducible physicochemical properties achieved through chemical functionalization, these resulting hydrogels offer the potential to create controlled, consistent, and relevant three-dimensional (3D) tissue models [38,39]. However, only a limited number of studies have sought to modify liver ECM hydrogels in order to enhance their physicochemical properties [41–43]. In our study, due to presence of collagen as the abandon protein in synthesized dECM, -NH2 chemical groups on the backbone of collagen structure are suitable to be modified and methacrylate groups could be substituted. For preparing the dECM for methacrylation process, the collagen should go through solubilizing process.

In this stage, pepsin has been utilized to enzymatically digest the matrix [44,45]. The action of pepsin involves cleaving peptide bonds at specific locations within the non-helical region of the collagen chain. This process effectively eliminates the non-helical immunogenic region of collagen, contributing to the removal of potentially immunogenic components from the matrix [46]. TPC of functionalized dECM was evaluated as the final product to be used in the bioink. It is shown that the TPC has been reduced to 55.95± 2.65 mg/g of dry LdECMMA sample (**Figure 1(D)**). To indicate the degree of substitution (DS) of methacrylate group, ^1^H NMR was done as presented in **Figure 1(E)**. The methacrylation of solubilized LdECM led to an appearance of new peaks at 5.42, and 5.77 ppm and at 1.94 ppm that can be attributed to the new methacrylate and methacrylamide end groups of dECM macromolecules specified **Figure 1(E)**. Integrating the peaks regarding methacrylate groups indicates around 56% of DS.

### 3.3 Measurements of mechanical, swelling and degradation properties

It has been proven that the mechanical properties of the fabricated scaffold have significant influence on the cell fate and differentiation, and also ECM remodeling [47,48]. Thus, we have measured the compression modulus of the hybrid GelMA-LdECMMA to investigate the influence of the LdECMMA concentration. By adding LdECMMA to GelMA, an improvement in mechanical properties was expected due to higher degree of crosslinking, as increasing the methacryloyl functional groups. In the hybrid mode, these are not only the GelMA polymer branches that make covalent connections during the photocrosslinking process, however, because of the presence of the C=C groups on the LdECMMA backbone, they will participate in the crosslinking, resulting in increasing internal connectivity. As the concentration of LdECMMA increased from 0 to 2% (w/v), a significant rise in the measured compression modulus was observed according to **Figure 2(A)**. The compression modulus of control GelMA was 2.06 ± 0.27 kPa and by adding 0.5%, 1% and 2% of LdECMMA, the compression modulus ended up to 6.02 ± 0.95, 13.19 ± 2.84 and 36.21 ± 10.12 kPa respectively. The mentioned results are showing that we can increase the compression modulus up to 1658% based on the control GelMA. This tunability in the mechanical properties makes the developed bioink as a promising biomaterial for different liver tissues modeling. Various studies have investigated the mechanical properties of healthy liver and different stages of diseased or cancerous liver [49,50]. The elastic modulus of the native and healthy human liver is reported to be around 12 kPa [51] which indicates that the hybrid bioinks containing 0.5% and 1% (w/v) LdECMMA are promising biomaterials to model the healthy liver ECM. However, during different stages of fatty liver disease toward cancerous liver, the stiffness of the liver tissue would increase due to fat accumulation. In this case, higher concentrations of the LdECMMA leading to higher mechanical properties would be the choice to represent the desired scaffold stiffness in liver disease in vitro modeling.

The swelling ratio of hydrogels plays a crucial role in tissue engineering as it impacts several factors such as surface properties, and solute diffusion [52]. By measuring the swelling ratio of crosslinked hydrogel samples, we can gain insights into the crosslinking degree and the ability of the hydrogel to absorb water [53]. This capability is influenced by the pore size of the scaffold as well as the interactions between the solvent and polymer. **Figure 2(B)** shows the swelling ratio data which by increasing the LdECMMA concentration, it is illustrating the reverse trend in comparison to the mechanical properties. The control GelMA sample is showing highest swelling ratio equal to 2302.64 ± 47.59%. Increasing the concentration of LdECMMA leads to a higher crosslinking density, resulting in a decrease in water retention capability. Consequently, bioinks containing LdECMMA exhibit significantly lower swelling ratios, with the swelling ratio for 2% LdECMMA reaching 1370.3 ± 94.6%, which is approximately 1.6 times lower than the control GelMA. Interestingly, the reduction in swelling ratio between 1% and 2% LdECMMA is not significant, suggesting that the water retention capability reaches a saturation point. Consequently, further increasing the LdECMMA concentration has a relatively insignificant effect on the swelling ratio. This finding indicates that there may be a limit to the impact of LdECMMA concentration on the water retention properties of the hydrogels.

To further investigate the mechanical properties and swelling behavior of the hydrogels, the microstructures were examined using the SEM. The SEM images, as depicted in **Figure 2(E)**, reveal that the GelMA hydrogel contains pores with a size range of 70 to 130 μm (with an average of 100.34 ± 27.99 μm). This pore structure contributes to lower mechanical stiffness and a higher capacity to absorb water, as discussed earlier. However, with the addition of LdECMMA to the bioink, there was a significant reduction in pore size. The hydrogel containing 2% LdECMMA exhibited a pore size of 29.13 ± 8.15 μm, which correlated with the highest compression modulus and the lowest swelling ratio. It is worth noting that the pore sizes observed in the LdECMMA-containing hydrogels did not show significant differences among themselves (**Figure 2(C)**). This observation suggests that there was a saturation point in the decrease of pore size, which aligns with the trend in swelling ratio discussed earlier. In summary, the SEM analysis revealed that the addition of LdECMMA to the hydrogel formulation led to a reduction in pore size, resulting in improved mechanical stiffness and reduced swelling ratio. This indicates that the incorporation of LdECMMA influenced the microstructure of the hydrogel, leading to enhanced controllability of mechanical properties significantly without meaningful influence on swelling behavior adversely.

Biodegradability has been a highly desirable characteristic in tissue engineering applications, particularly for hydrogel materials, as it indicates how long the scaffold can maintain its structural integrity. However, hydrogels typically exhibit slow degradation rates when placed in PBS solution without any enzymes, requiring a significant amount of time for degradation to occur. To investigate the degradation properties of GelMA-LdECMMA hydrogels, a series of experiments were conducted. Firstly, the dry weight of the samples was measured, and they were then rehydrated. Next, hydrogels with varying compositions were immersed in a collagenase solution and incubated at 37 °C. The weight of the hydrogels was recorded at different time intervals after lyophilization. As demonstrated in **Figure 2(D)**, the GelMA control sample was found to be completely degraded within less than 4 hours. In contrast, the hybrid bioinks containing 1% and 2% LdECMMA exhibited a significantly slower degradation rate, preserving their structure for over 2 days. This outcome clearly indicates that the incorporation of LdECMMA extended the degradation time of GelMA. The differences in degradation rates between GelMA and the hybrid hydrogels can be attributed to two main factors. Firstly, the introduction of an additional methacrylated component to GelMA resulted in a higher degree of crosslinking. Consequently, during the degradation process, a greater number of covalent bonds needed to be broken down to cause structural collapse. Additionally, through swelling and microstructural evaluations, it was observed that higher concentrations of LdECMMA led to a reduction in pore size within the hydrogels.

Consequently, during enzymatic degradation, the hydrogels absorbed a smaller amount of the solution, resulting in a decrease in the rate of degradation [54]. The ability to tune the degradation rate of these hybrid hydrogels makes them highly promising biomaterials for a wide range of applications, both in vivo, such as for implantations, and also long-term in vitro application [55], including drug screening devices.

Furthermore, structural analysis of samples containing dECM revealed the presence of wrinkles in the GelMA walls, as depicted in **Figure 2(E)**. These wrinkles became more pronounced with an increased concentration of LdECMMA, suggesting a direct correlation. This phenomenon can be attributed to the intrinsic structural characteristics of LdECM and LdECMMA, as similar wrinkled structures are also observed in their SEM images (**Figure S4**).

To validate the impact of dECM methacrylation on the physiological characteristics of the engineered hydrogels, control prepolymer samples incorporating unmodified dECM were prepared for comparison with the existing findings. The incorporation of various concentrations of solubilized dECM into 5% GelMA resulted in an enhancement of mechanical properties, likely attributed to the increased overall concentration of the prepolymer solution, as illustrated in **Figure S5(A)**. Nonetheless, the magnitude of this increase did not match that observed in samples containing LdECMMA. The integration of LdECMMA into GelMA, followed by light-induced crosslinking, caused additional chemical covalent bonds not only among GelMA chains but also between GelMA and LdECMMA fibers, unlike in samples with unmodified dECM where covalent bonding occurred solely among GelMA chains, excluding dECM fibers. Furthermore, in samples with unmodified dECM, the absence of any bonding between dECM fibers and the 3D structure of GelMA led to a significantly faster degradation by collagenase enzyme in the enzymatic degradation assay, particularly evident in the 5G0.5dECM samples compared to both 5G0d and 5G0.5d samples, as shown in **Figure S5(B)**.

### 3.4 Crosslinking Time Measurements and Rheological Characterization

The effective utilizing of dECM-based bioink in SLA bioprinting relies on a material that boasts rheological properties that are finely tuned. To achieve this, an assessment of both the viscosity and the crosslinking kinetics was conducted via rheological characterization. Illustrated in **Figure 3** are the viscosity and dynamic moduli for various formulations of dECM-based bioinks. As depicted in **Figure 3(A)**, the incorporation of LdECMMA into GelMA significantly enhances the viscosity of the bioinks relative to the control sample, a consequence of the rich presence of collagen fibers in LdECMMA. It has been established in literature that the viscosity of the resin for optimal usage in SLA 3D printing should not surpass 5000 cPs [56,57]. Thus, this suggests that bioink formulations containing 2% LdECMMA may not be ideally suited for SLA bioprinting applications. However, owing to the pronounced shear-thinning property exhibited by the 5G2d sample, it emerges as a promising candidate for extrusion bioprinting, as verified by various studies exploring the extrudability of dECM-based bioinks in 3D bioprinting contexts [58,59].

Furthermore, the photorheological properties of the bioink were elucidated using a photorheological test, measuring the storage (G’) and loss (G”) modulus of diverse hybrid inks over 60 seconds in the dark to establish a baseline, followed by illumination with visible light to monitor changes in the rheological properties of the ink. **Figure 3(B)** displays the evolution of G’ and G” during the photocuring process, offering insights into several critical aspects. The point at which the curves of G′ and G″ intersect is typically regarded as the material’s gel point [60]. In the case of the 5G0d sample, this gel point occurs approximately 30 seconds after the commencement of irradiation (**Table S1**). Conversely, in samples containing LdECMMA, the gel point does not occur within the photocuring timeframe, as indicated by the G’ values exceeding those of G” prior to irradiation, likely due to presence of collagen fibers that through entanglement, partial network has been formed prior to photo-gelation which is more probable in high viscosity inks. Additionally, the period from the onset of exposure to the observable rise in G’ and the initiation of gelation, known as gel time (GT), varies across samples [60]. For the 5G0d sample, gelation starts after a delay of about 20 seconds (**Table S1**), whereas, in samples with a high concentration of methacryloyl groups, the photo-crosslinking reaction initiates instantly upon light exposure, as evidenced by the absence of delay in the rheological data.

A crucial consideration in SLA printing is the ability to cleanly separate the printed structure from the uncured bioink, which is pivotal for achieving high printing resolution. Optimal printing necessitates a printed structure with high stiffness and an uncured bioink with low viscosity. ΔG’ is quantified as the logarithmic difference between the G′ values of the crosslinked and uncrosslinked states, serving to articulate the rheological disparity between the printed structure and the uncured hydrogel. Throughout the photocuring process, as the bioink undergoes increased crosslinking, the ΔG′ value rises until crosslinking is complete, whereupon it stabilizes [60]. Through this experiment, samples were illuminated for 5 minutes to ensure that G’ reached its saturation point, signaling the completion of photocuring. The variations in mechanical properties previously discussed are also reflected in different final G’ and G″ values after crosslinking, which rise with increasing LdECMMA concentration, thereby elevating ΔG′ as well (**Table S1**). Moreover, literature indicates that achieving the optimal crosslinking density during photo-crosslinking is facilitated by aiming for 75%–80% of the ΔG’ logarithmic value, defining this range as the printable window [60]. **Table S1** presents the printable windows for various bioinks, illustrating that an increase in LdECMMA concentration shortens the crosslinking time, attributed to a higher concentration of methacryloyl groups and a reduced pathway for the diffusion of free radicals.

Lastly, sustaining the printed structure’s shape requires greater elasticity than viscosity. The difference between G′ and G″ values, the larger it is, the better the shape retention. The term ΔM denotes the logarithmic difference between the G′ and G″ values following full exposure during photocuring [60]. **Table S1** details ΔM values for different bioinks, revealing that as LdECMMA concentration rises, so does the degree of crosslinking, yielding a stiffer structure and, consequently, a larger ΔM value.

To explore the viscoelastic properties of crosslinked hydrogels, **Figure S6** presents the comparison between the G’ and G” of photocrosslinked hydrogels across various angular frequencies. Throughout the entire range of angular frequencies tested, the G’ for all hydrogel formulations consistently exceeded G’’, indicating a gel-like viscoelastic behavior. For hydrogels incorporating LdECMMA, G’ and G’’ moduli exhibited frequency independent behavior over the entire range of the angular frequency tested, suggesting gel-like behavior. Conversely, the 5G0d sample exhibited a deviation from this pattern, with G’ and G’’ not maintaining independency behavior throughout the angular frequency range, reflecting its unstable structural integrity [61,62].

### 3.5 Printability evaluation

As discussed in existing literature, an optimal photocurable bioink must meet specific criteria, including rapid photo-crosslinking, high crosslinking density, and enhanced fluidity. Firstly, a high crosslinking rate is imperative to expedite curing, thus reducing printing time and enhancing printing resolution. Secondly, maintaining a high crosslinking density is essential to impart adequate stiffness, thereby preventing deformation. Lastly, high fluidity is critical for improving printing efficiency, facilitating rapid flow of uncured hydrogel bioinks into gaps and spaces, and simplifying the removal of uncrosslinked ink to achieve the desired final shape [63].

To assess the printability of various compositions of GelMA and LdECMMA inks, a pattern featuring seven hexagons was subjected to different durations of visible light irradiation for crosslinking. **Figure 4(A)** illustrates the printed structures corresponding to these compositions. The control sample, 5G0d, required a minimum irradiation time of 4 minutes for crosslinking. Subsequently, exposure for 4 minutes yielded a stable structure with undulating and wavy walls, while a longer exposure of 5 minutes resulted in increased stiffness of the walls. Conversely, presence of 0.5% LdECMMA led to the formation of a stable and rigid structure after just 1 minute of visible light exposure. However, after 3 minutes of photocrosslinking, significant over-crosslinking occurred, leading to noticeable reduction in the size of holes within the hexagons. Similarly, for the 5G1d sample, a 30 second exposure produced a moderately stable structure with distinguishable but broken branches, while a 1 minute exposure yielded a firm and stable structure. However, exposure exceeding 1 minute led to over-crosslinking of the 5G1d ink. Furthermore, increasing the LdECMMA content to 2% resulted in incomplete patterning even after a 30 second exposure, with over-crosslinking observed after 1 minute of irradiation. Overall, increasing the LdECMMA content enhanced the speed of crosslinking by enhancing the methacrylol groups, thereby enabling the formation of stable structures with reduced exposure time. Consequently, the dECM-based bioink fulfilled the criteria for rapid photocrosslinking and high crosslinking density.

**Figure 4.**
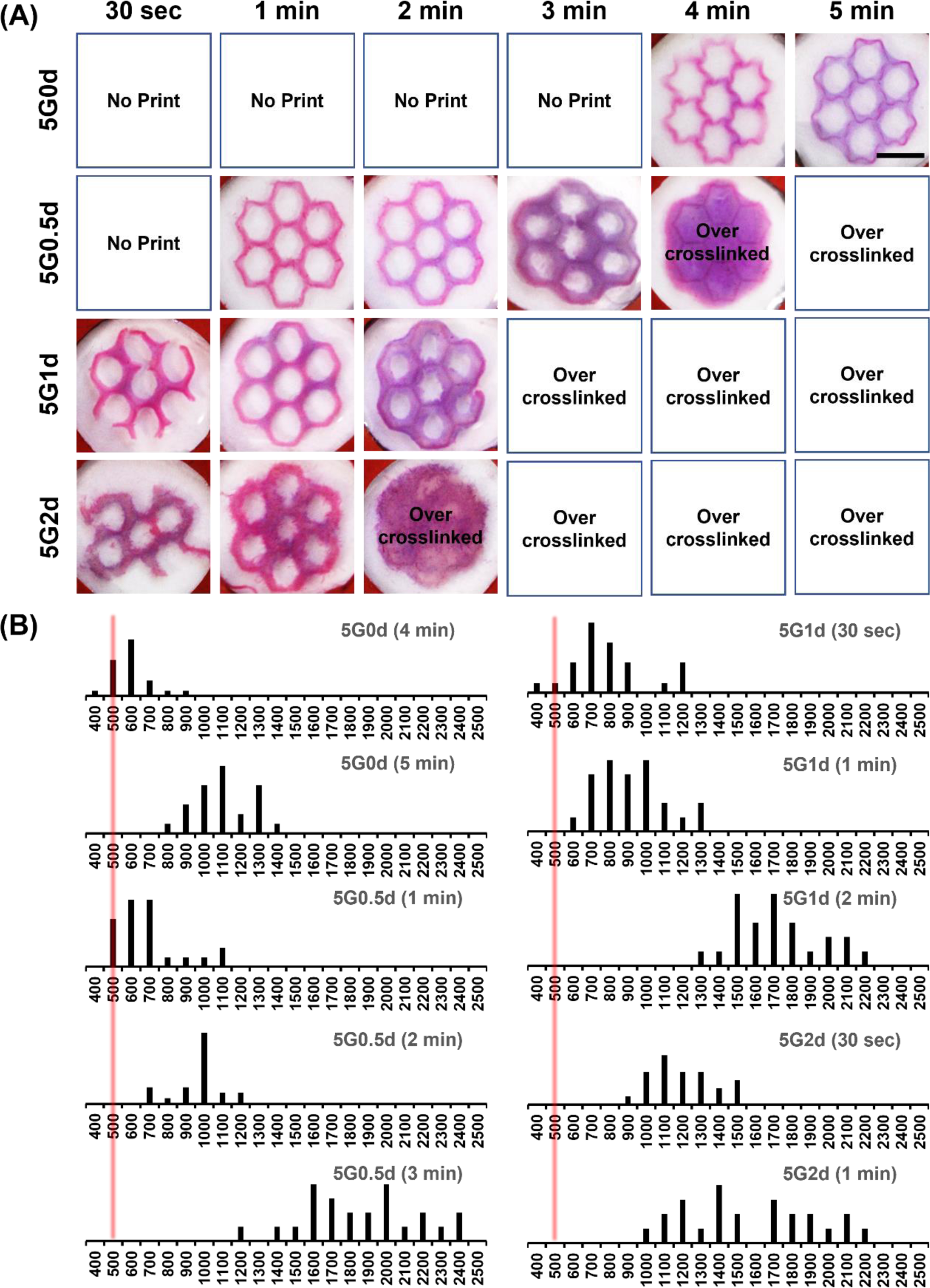
SLA printability evaluation. (A) Photo-patterning of different compositions of LdECMMA-based hydrogels under various exposure times, The fabricated scaffolds were stained with red dye for better visualization (scale 5 mm). (B) Printed line thickness measurement of different constructed patterns.

In the evaluation of suitable inks for SLA printing, fluidity emerges as a crucial third criterion as mentioned previously. Rheological examinations, as discussed in the preceding section, have demonstrated that all examined inks, with the exception of the 5G2d ink, fall within this acceptable viscosity range. Notably, the viscosity of the 5G1d ink approaches the upper limit of this range. Despite the fact that both the 5G1d and 5G2d inks could be crosslinked within a 30-second timeframe, their relatively higher viscosity presented substantial difficulties. Specifically, the process of washing away uncrosslinked ink proved to be particularly problematic for these samples, leading to the occurrence of branch breakage post-printing.

Nevertheless, the photo patterning duration observed diverges from the optimal crosslinking time outlined in the prior section (**Table S2**). This difference can be attributed to the varying sensitivities of rheology and photo patterning investigations towards alterations in the crosslinking density and the architecture of the network. Rheological analysis is concerned with the assessment of the overall mechanical properties of the bulk material, in contrast to photo patterning studies which might prioritize the crosslinking at the surface or within specific localized areas. Moreover, photo patterning could involve the utilization of samples that are thinner or prepared under different conditions. The thickness of the sample, its shape, or the manner in which it is exposed to light are factors that can significantly impact the kinetics of crosslinking, potentially resulting in variations in the determination of optimal crosslinking times across these methodologies.

Resolution is a critical factor in SLA bioprinting that needs careful consideration. The resolution of printing is fundamentally influenced by the rate and density of photo-crosslinking, which are, in turn, dictated by the production rate and concentration of free radicals [60]. A more rapid photo-crosslinking process minimizes the opportunity for monomers or oligomers to diffuse from the intended area, thereby ensuring more precise control over polymerization sites. This precision facilitates the creation of finer features and more defined boundaries, leading to an enhancement in resolution [64,65].

To assess resolution, two key parameters are examined: the thickness of printed lines and the clarity of the edges of hexagonal structures which is reported in **Figure 4(B)**. In terms of line thickness, the control sample, designated as 5G0d, displayed a branch thickness of approximately 1054.16 ± 159.32 μm following a 5-minute crosslinking period. In contrast, the complete structure printed using 5G0.5d ink after just 1 minute of photo-patterning exhibited line thicknesses of 723.88 ± 169.21 μm, which is comparable to the thickness of lines printed with 5G0d ink over 4 minutes, measuring around 556 ± 102.19 μm, although with wavy lines. Additionally, the 5G0d ink after 5 minutes yielded relatively sharp outer edges, despite the areas within the hexagons appearing somewhat circular. Conversely, structures printed with 5G0.5d ink after 1 minute of crosslinking showcased sharpness in both outer and most inner edges. This comparison indicates that the incorporation of 0.5% LdECMMA into 5% GelMA not only significantly reduces the crosslinking time by a factor of five but also enhances the resolution. It is important to note, however, that adding LdECMMA to GelMA results in reduced transparency, leading to increased light scattering and, consequently, a potential decrease in printing resolution [66,67]. This phenomenon might explain the observed increase in branch thickness in structures printed with 5G1d ink after 1 minute of crosslinking, which was approximately 868 ± 181.48 μm. Despite these challenges, the outer edges of hexagons printed in this experiment remained sharply defined and well-formed.

Finally, for the 5G2d ink, the substantial decrease in transparency coupled with a significant increase in viscosity led to an increase in line thickness to 1143.21 ± 182.88 μm and 1518 ± 349.43 μm after 30 seconds and 1 minute of photo-patterning, respectively. Moreover, the high line thickness resulted in the absence of sharp edges, rendering this ink less suitable for SLA printing applications due to its compromised resolution.

### 3.6 Viability evaluation and metabolic assay

To explore the viability of HepG2 cells encapsulated in both control samples and those containing LdECMMA, bioprinted constructs underwent crosslinking and were subsequently cultured for approximately one month. During various stages of the culture period up to 14 days, live/dead staining was performed on the scaffolds to assess cell viability. The imagery presented in **Figure 5(A)** illustrates that, across all time points and in both environments, the majority of cells not only endured the printing process but also proliferated within the scaffolds. Previous research has noted that HepG2 cells tend to aggregate into clusters as time progresses, with these clusters growing in size and eventually merging to form larger colonies [68–70]. A distinct difference was observed between cells in the 5G0d and the 5G0.5d scaffolds, with cells in the latter tending to cluster more rapidly than in the control. Given the cellular tendency for cluster formation, traditional individual cell counting methods were inadequate for viability assessment.

**Figure 5.**
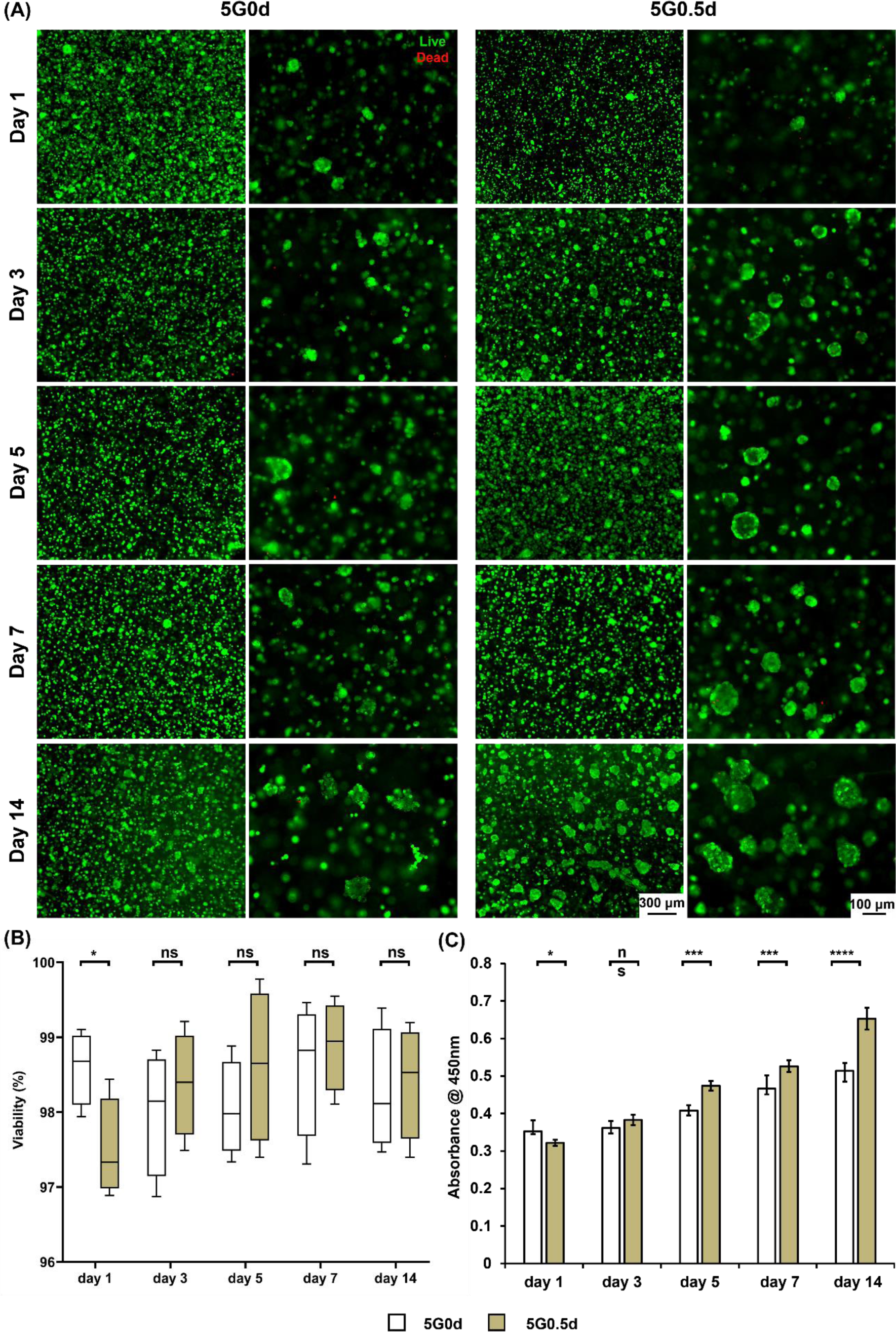
Biocompatibility and metabolic characterization of LdECMMA containing hydrogel. (A) Representative images showing live and dead HepG2 cells encapsulated in the hydrogels over 14 days of culture. (B) Cell viability evaluation calculated by the number of the live and dead cells (n=3). (C) XTT assay showing HepG2 cells metabolism and proliferation within 14 days of culturing cells in 3D hydrogels structure (n=5). (* p < 0.05, ** p < 0.01, *** p < 0.001 and **** p < 0.0001)

Instead, cell viability was quantified based on the surface area ratio of live to dead cells. Throughout the 14 days culture period, the viability of HepG2 cells remained above 95% in both scaffold types, indicating the biocompatibility of both GelMA and GelMA enhanced with LdECMMA (**Figure 5(B)**). Initially, cell growth was more pronounced within the 5G0d scaffold compared to its LdECMMA-supplemented counterpart, likely due to the former’s lower mechanical properties and higher swelling ratio [71]. However, from day 3 onwards, the incorporation of LdECMMA rendered the scaffold more alike to native liver tissue, thereby fostering enhanced proliferation in the 5G0.5d samples. From days 3, 5, 7, and 14 of culture, cell clusters in the 5G0.5d samples exhibited accelerated growth in both size and quantity, as evidenced by live/dead imaging data in Figure 3A. In addition to visual assessments, an XTT assay was conducted to measure the metabolic activity of the cells, further quantifying cell proliferation (**Figure 5(C)**). Initially, the 5G0d scaffold housed more cells compared to the 5G0.5d sample, resulting in higher absorbance readings due to the control hydrogel’s lower stiffness. Nevertheless, as HepG2 cells within the LdECMMA-enriched scaffold proliferated from the third day of culture, they experienced an environment more closely resembling the native liver ECM niche, which in turn significantly enhanced their metabolic activity compared to those in the 5G0d scaffold. This suggests a promising capability of LdECMMA-based scaffolds to support liver tissue regeneration.

### 3.8 Cell proliferation and morphology assessment

To examine the cellular morphology and organizational patterns within the bioprinted hydrogel scaffolds, staining of the cytoskeleton and nuclei was conducted at various culture intervals, as depicted in **Figure 6**. Utilizing GelMA, recognized for its cytocompatibility and superior photo-patterning capabilities, was enriched with LdECMMA, known for its abundance in intricate extracellular matrix proteins and growth factors.

**Figure 6.**
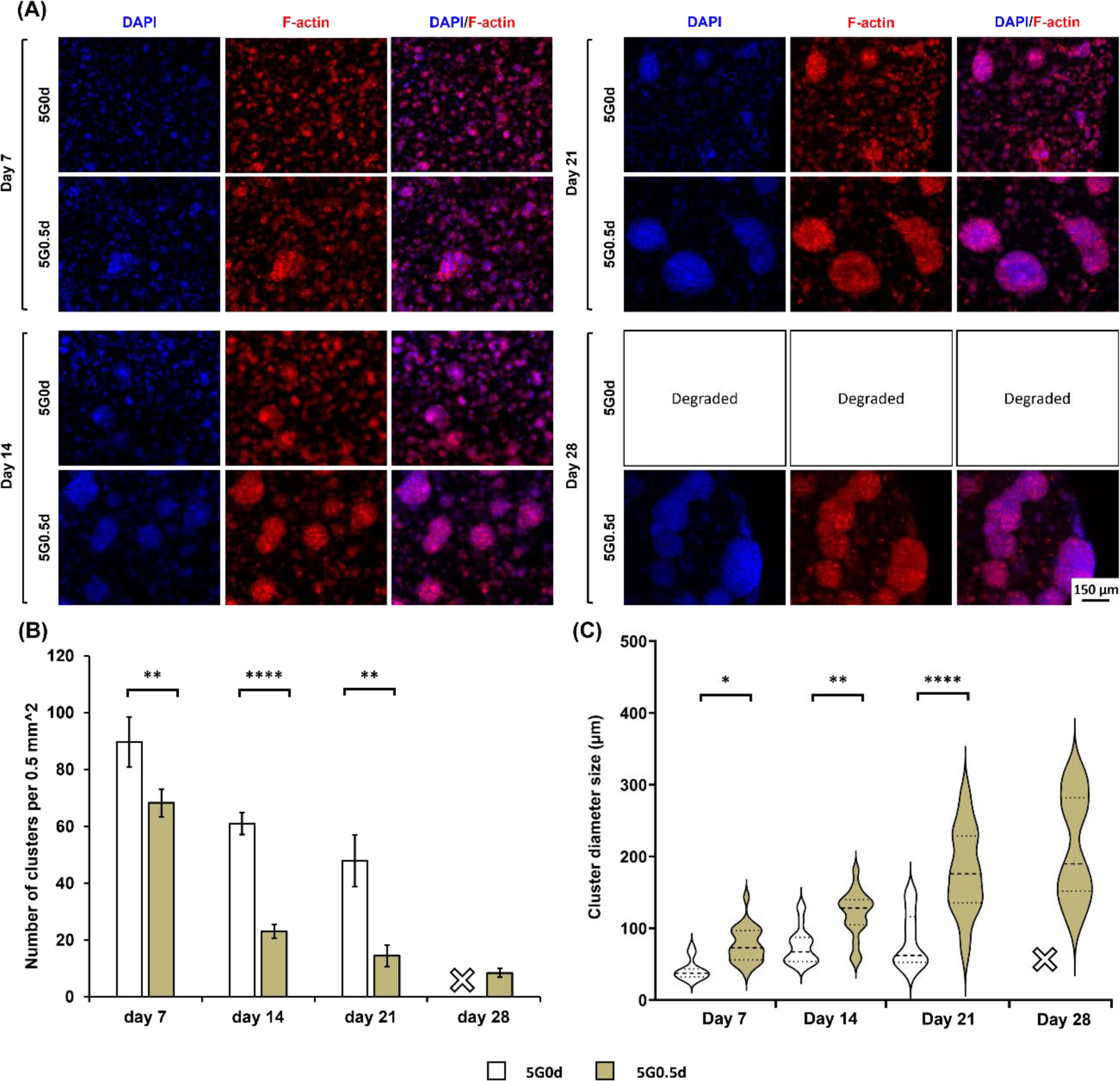
Evaluating cell growth, proliferation and morphology over 28 days of culture. (A) Fluorescent images showing HepG2 cells morphology during the culture period with phalloidin stained F-actin (red) and DAPI stained nuclei (blue) and the merged pictures (scale 150 µm). (B) HepG2 cell cluster number per unit of area on different days of the culture (n=5). (C) HepG2 cluster size distribution over different days of the culture (n=5). (* p < 0.05, ** p < 0.01, *** p < 0.001 and **** p < 0.0001)

The findings demonstrated a progressive development of HepG2 cells, evolving from individual cells into cohesive clusters, and ultimately forming multicellular spheroids, similar to studies reported previously [72,73]. These spheroids remained viable and stable for the duration of the study, exhibiting uniform distribution across the three-dimensional scaffold structure. In both the control and the LdECMMA containing bioink samples, cluster formation was evident by day 7 (**Figure 6**), with **Figure 6(B)** and **(C)** showcasing the number of clusters per unit area and the average cluster size, respectively. On day 7, the LdECMMA enriched samples (5G0.5d) not only exhibited fewer clusters than the control samples (5G0d) but also displayed significant differences in cluster sizes, 76.86 ± 24.73 μm for 5G0.5d samples versus 41.76 ± 14.11 μm for the control, indicating more pronounced cell proliferation in the LdECMMA samples even within the first week of culture. By day 14, clusters in the LdECMMA supplemented GelMA were notably larger and more distinct compared to those in 5G0d, emphasizing the enhanced cellular proliferation.

Clusters in the presence of LdECMMA demonstrated a more coherent and structured arrangement, characterized by a well-defined actin filament network and multicellular aggregates that expanded in size as the culture period progressed, in contrast to the clusters formed in the GelMA-only hydrogels. By the third week of culture, the average cluster size in the 5G0d and 5G0.5d samples had grown to 179.27 ± 59.87 μm and 82.98 ± 34.00 μm, respectively (**Figure 6(C)**). Additionally, a decreasing trend in cluster number was observed for both sample types, attributed to the merging of clusters. As highlighted in the degradation study, the 5G0d samples exhibited the highest rate of degradation and, compounded by the cellular presence, did not maintain structural integrity until day 28 of culture and was completely degraded. Conversely, **Figure S7** illustrates that in the 5G0.5d samples, clusters began merging from day 21, leading to the formation of larger cell aggregates by day 28, occupying significant portions of the gel structure.

In summary, the incorporation of LdECMMA into the hydrogel scaffolds significantly promoted HepG2 cell spreading and proliferation. This enhancement is likely facilitated by the extracellular matrix proteins and growth factors inherent to the dECM component within the scaffold, and potentially through modifications to the mechanical strength and architecture of the GelMA network [72].

### 3.8 Albumin secretion analysis and immunostaining

To verify the preservation of HepG2 cell functionality within the hydrogel scaffolds, the production of albumin—a marker of hepatocyte function—was assessed through immunostaining at various culture intervals (days 7, 14, 21, and 28) [74,75]. Evidence of albumin secretion was detected at all evaluated time points in both experimental setups, affirming the functional capacity of the bioprinted HepG2 cells within the hydrogels. Notably, HepG2 cells exhibited a marked increase in albumin production over time, particularly when cultured in hydrogels enriched with LdECMMA, in comparison to those in the control hydrogel. As illustrated in **Figure 7(A)**, albumin secretion was predominantly localized within HepG2 cell clusters formed in the 3D scaffold structure. Despite both sample types exhibiting localized albumin production within cell aggregates throughout the culture duration, the control samples (5G0d) displayed a lack of significant albumin secretion in some nuclei clusters after the first and second weeks of culture. Conversely, the LdECMMA-enhanced hydrogels (5G0.5d) consistently showed significant albumin secretion across the majority of matured clusters. Moreover, a combined assessment of morphology and immunostaining for Albumin is presented in **Figure S8**, illustrating the integration of structural evaluation with specific protein expression analysis.

**Figure 7.**
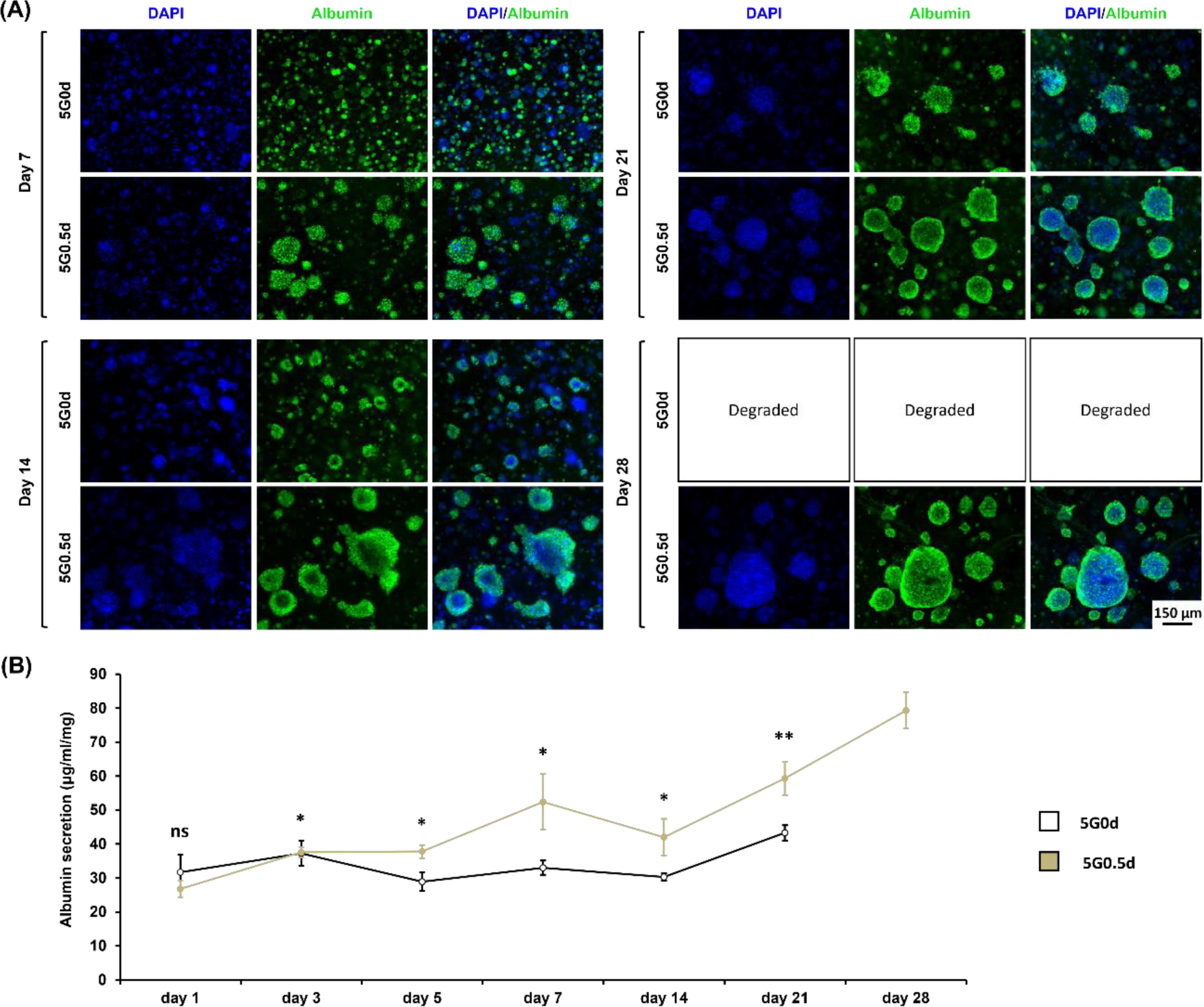
Liver functionality evaluation of 3D cultured HepG2 within hydrogels. (A) Albumin secretion activity visualized through immunostaining of different days of the culture (scale 150 µm). (B) Albumin production by HepG2 over the culture period in dECMMA containing hydrogel (n=5). (* p < 0.05, ** p < 0.01, *** p < 0.001 and **** p < 0.0001)

In addition to immunostaining, albumin secretion by HepG2 cells within the hydrogels was quantified using an ELISA kit, with the findings summarized in **Figure 7(B)**. From day 3 onwards, the 5G0.5d samples demonstrated significantly higher albumin secretion levels compared to the control samples, attributed to the enhanced cell proliferation observed in the LdECMMA-supplemented hydrogels. Specifically, albumin secretion levels increased from 31.69 ± 5.25 to 43.31 ± 2.25 μg/ml/mg (representing a 1.4-fold change) and from 26.79 ± 2.41 to 79.28 ± 5.31 μg/ml/mg (indicating a 2.9-fold change) for the 5G0d after day 21 and 5G0.5d after day 28 of culture, respectively. This progression underscores the significant impact of LdECMMA incorporation on enhancing HepG2 cell functionality within the scaffold environment.

### 3.9 Histology assessment

The objective of this section is to shed light on the histological variances observed in HepG2 cells cultured on GelMA scaffolds, both with and without the addition of LdECMMA. Utilizing a consistent H&E staining protocol across both sets of samples, we aimed to define differences in cellular morphology and the interplay with the matrix. In sections stained with H&E, HepG2 cells within the pure GelMA scaffolds (**Figure 8**) display the expected cytoplasmic staining characteristics, showcasing nuclei distinctly bordered by the stark blue of the hematoxylin and a cytoplasm that stands out in a contrasting lighter pink due to the eosin. In contrast, HepG2 cells within the GelMA scaffolds that were enriched with LdECMMA (**Figure 8**) are marked by a significantly more intense cytoplasmic staining. This suggests a unique interaction with the eosin dye, which is an acidic stain that preferentially associates with basic cellular components, predominantly the protein elements situated in the cytoplasm. Verifying with literature, the ECM is recognized for its imperative influence over cellular conduct and the modulation of gene expression, which sequentially impacts the synthesis of proteins within the cytoplasm [76–78]. Consequently, the deeper staining of the HepG2 cells’ cytoplasm in the LdECMMA-infused sample could be reflective of an enhanced synthesis of cytoplasmic proteins, potentially due to advanced cellular maturation processes.

**Figure 8.**
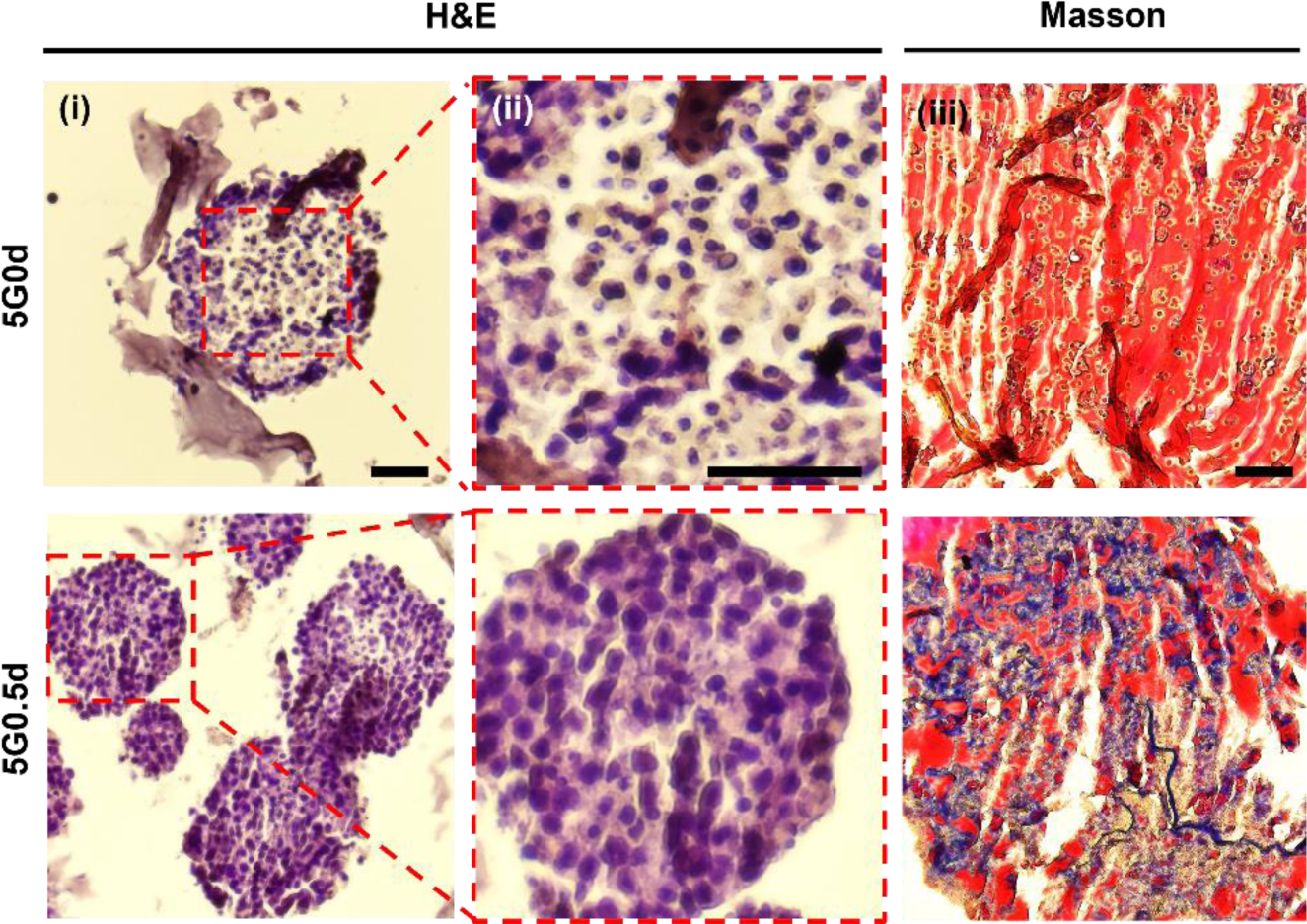
Histological evaluation of bioprinted structures. H&E and Masson Trichrome staining of control and LdECMMA containing samples after 28 days of HepG2 cells culture (scale 50 µm)

Furthermore, the presence of unoccupied spaces within the clusters of cells observed in the GelMA-only sample, as opposed to the densely populated cellular arrangement seen in the 5G0.5d sample, might stem from different causes. These include, but are not limited to, heightened cellular proliferation and migration, potentially spurred by the growth factors and other signaling molecules contained within the dECM-supplemented sample, which contribute to the reduction of voids within the cellular clusters. Additionally, Masson’s trichrome staining was employed to ascertain the presence of collagen fibers within the scaffolds utilized for HepG2 culture. In the 5G0d sample, as illustrated in **Figure 8**, an absence of collagen fibers is noted, correlating with the initial lack of collagen supplementation. Conversely, the 5G0.5d sample reveals that the collagen fibers, intrinsic to the LdECMMA, have been retained within the structure in around one month of culture, likely due to covalent linkages formed with the GelMA’s backbone, thus exhibiting lower degradation rates. This finding underscores the sustained incorporation of LdECMMA within the GelMA scaffold as being advantageous for the prolonged cultivation of liver cells.

## 4. Conclusion

The challenges posed by the inadequate mechanical and physical properties, along with the limited printability of dECM containing bioinks, have been significant barriers to their application in 3D bioprinting technologies. To address these issues, our research introduced a novel liver dECM-based bioink, created by methacrylating liver dECM. This innovation led to a hybrid LdECM-based hydrogel that demonstrates adjustable physicochemical properties and rapid crosslinking under visible light. Extensive testing confirmed that this new bioink significantly improves mechanical and rheological properties, as well as its capability for precise photo-patterning. Furthermore, in vitro experiments using HepG2 cells revealed outstanding biocompatibility, with evidence of enhanced cell proliferation and liver functionality compared to control samples. Looking ahead, the liver dECM-based bioink developed in this study holds promising potential as a sophisticated resource for regenerative medicine and drug screening endeavors.

## Acknowledgement

This work was supported by a Natural Sciences and Engineering Research Council of Canada (NSERC) Discovery Grant (RGPIN-2014-04010) and Canada Foundation for Innovation John R. Evans Leaders Opportunity Fund.

## Abbreviations

3D: three-dimensional
GelMA: gelatin methacrylate
ECM: Extracellular matrix
dECM: decellularized extracellular matrix
LdECMMA: human-on-a-chip
SDS: sodium dodecyl sulfate
DLP: digital light processing
EY: eosin Y
TEOA: triethanolamine
SEM: scanning electron microscopy
PBS: phosphate buffered saline
H&E: hematoxylin and eosin

