## Supplementary file for "3D bioprinting of Liver Microenvironment Model Using Photocrosslinkable Decellularized Extracellular Matrix based Hydrogel"

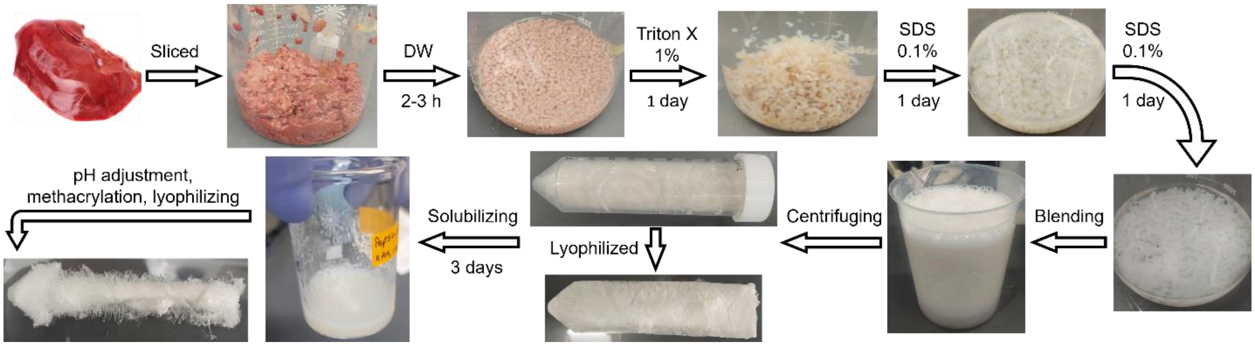


**Figure S1** LdECMMA synthesis process

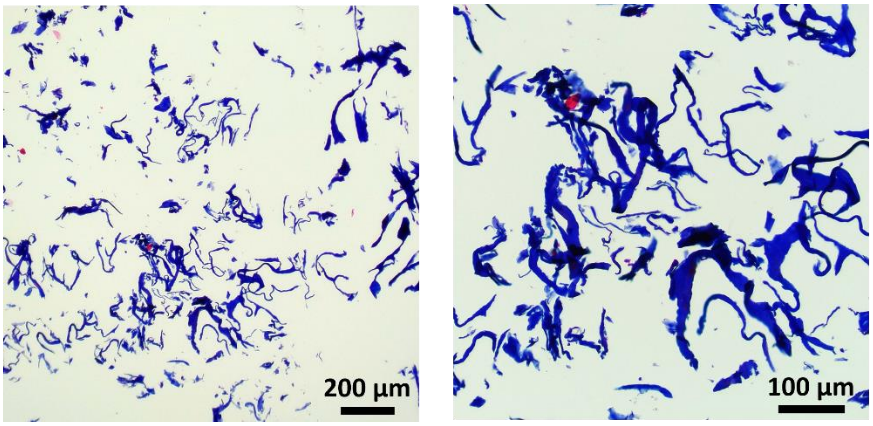


**Figure S2** LdECMMA masson staining for collagen fibers

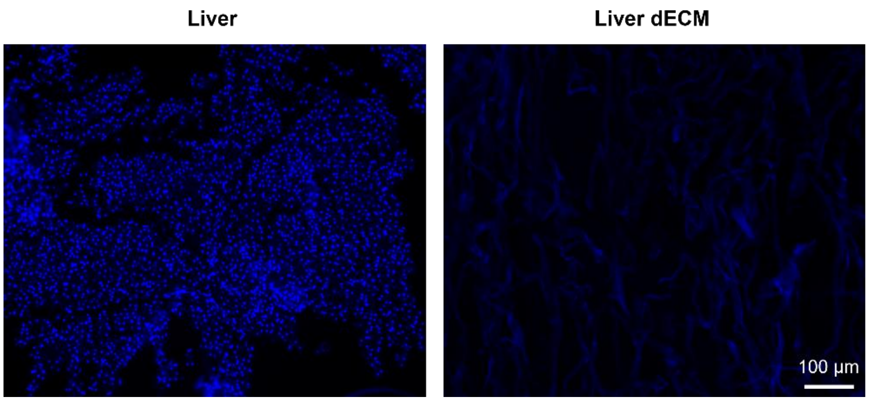


**Figure S3** Native liver and LdECM DAPI staining

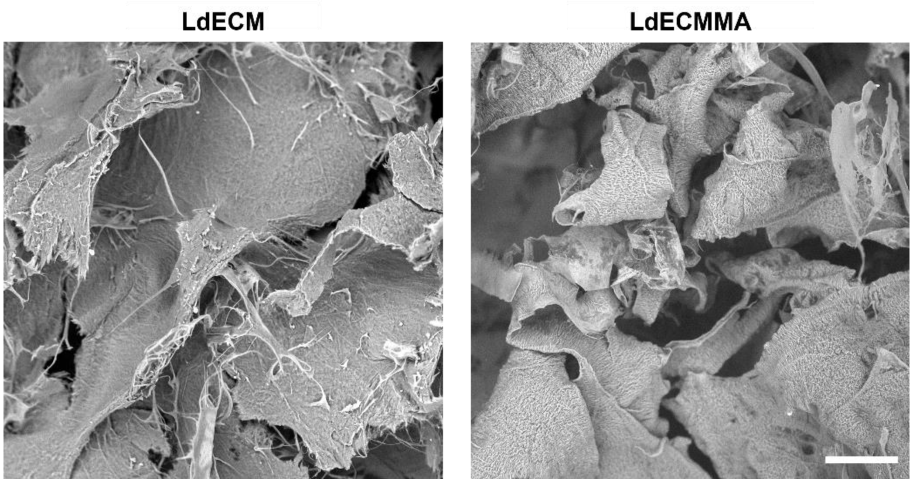


**Figure S4** LdECM and LdECMMA SEM imaging

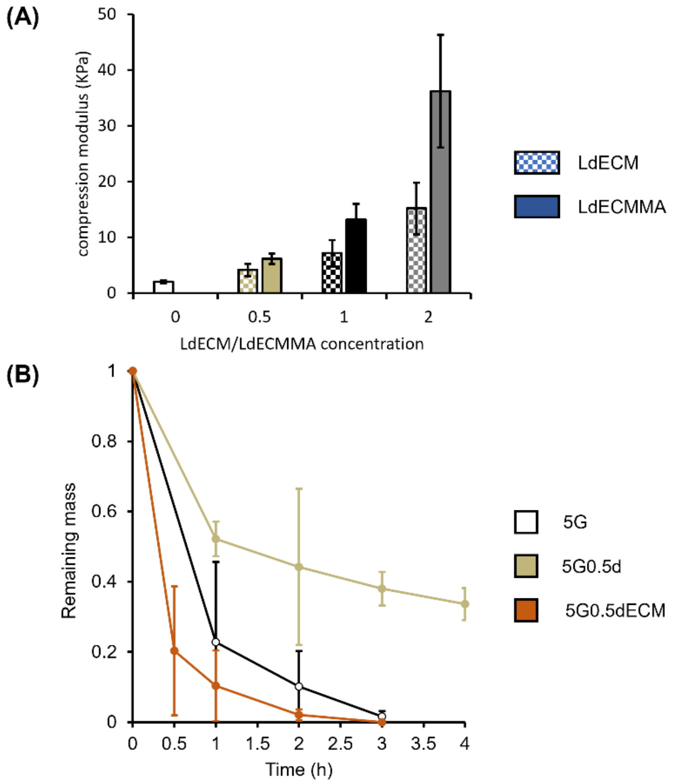


**Figure S5** dECM based hydrogels properties evaluation, (A) GelMA-LdECM hydrogels mechanical properties, and (B) degradation behavior comparison of LdECM and LdECMMA based hydrogel

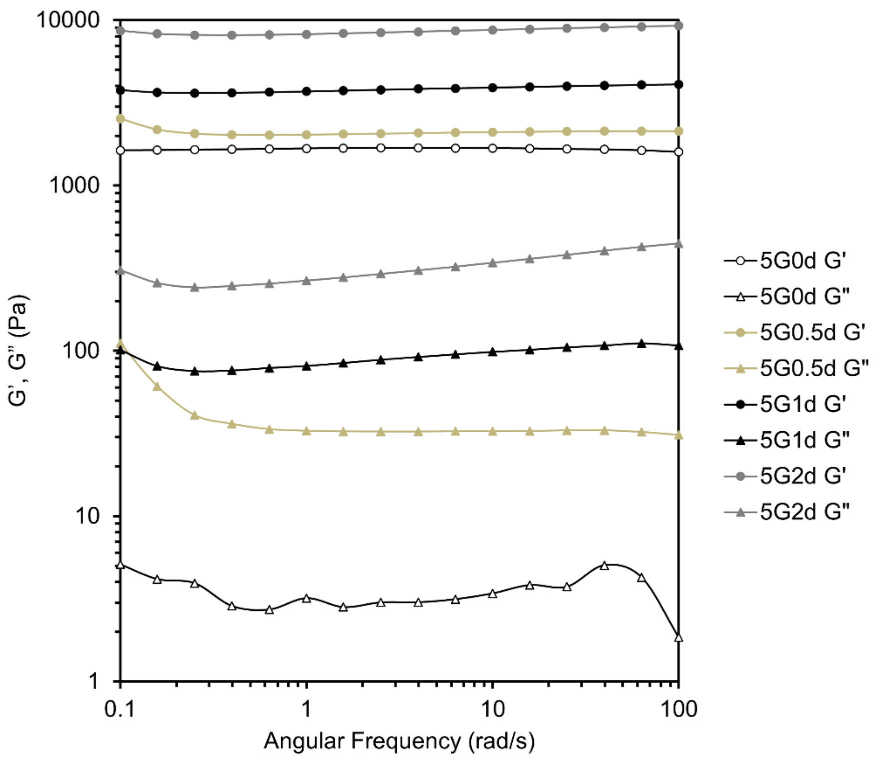


**Figure S6** Angular frequency swap evaluation of the different compositions

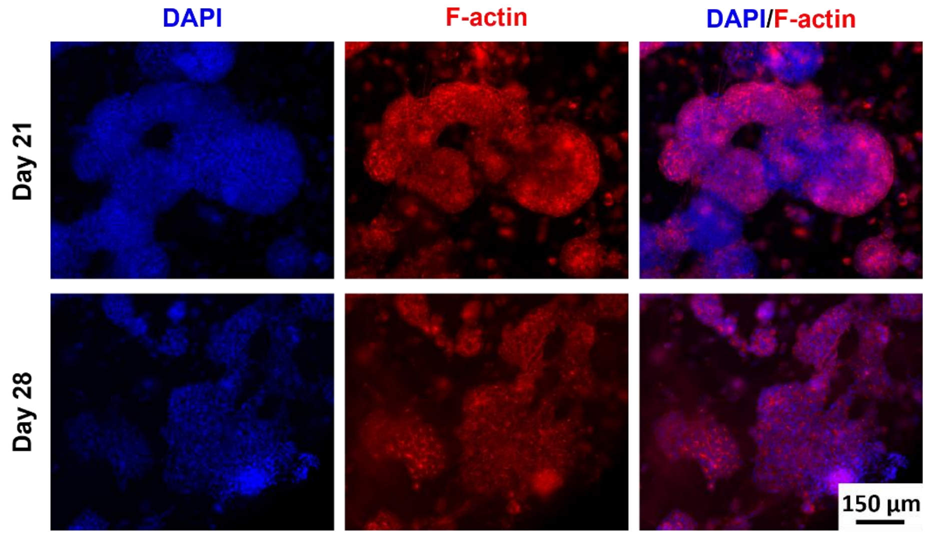


**Figure S7** Morphology evaluation of the HepG2 cells encapsulated in 5G0.5d hydrogel after 21 and 28 days of the culture showing cell clusters merging

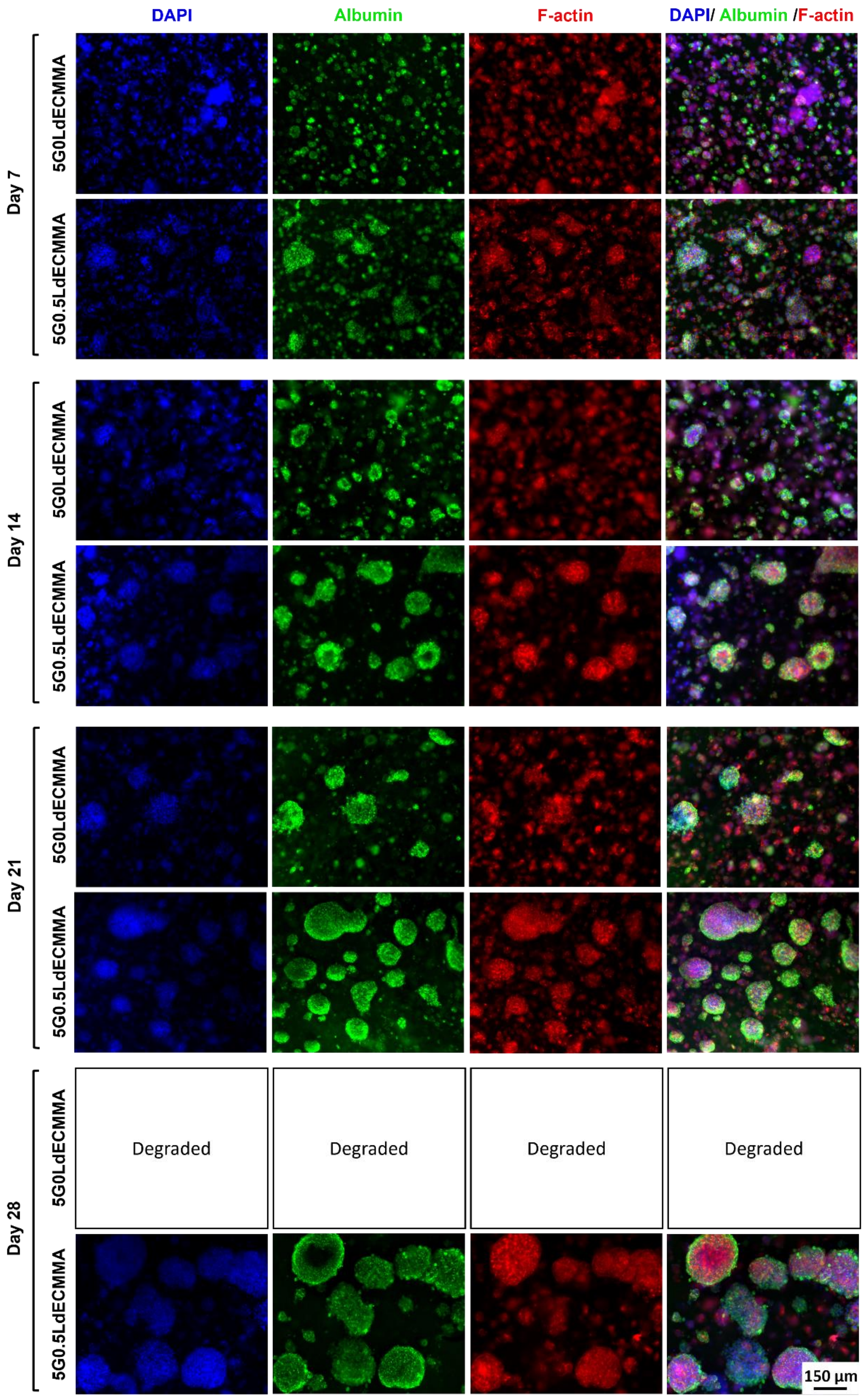

DAPI
Albumin
F-actin
DAPI/ Albumin /F-actin
Day 7
5G0LdECMMA
5G0.5LdECMMA
Day 14
5G0LdECMMA
5G0.5LdECMMA
Day 21
5G0LdECMMA
5G0.5LdECMMA
Day 28
5G0LdECMMA
Degraded
Degraded
Degraded
Degraded
5G0.5LdECMMA
150 µm

**Figure S8** Immunohistochemistary and morphological evaluation of the 3D cultured samples

Table S1

| *sample* | Gel time (s) | Gel point (s) | ΔG' (Pa) | Printable window (min) | ΔM (Pa) |
|---|---|---|---|---|---|
| *5G0d* | ≈ 24 | ≈ 30 | 1593.40 | 4.3-4.6 | 1589.775 |
| 5G0.5d | 0 | - | 2067.89779 | 4-4.2 | 2036.051 |
| 5G1d | 0 | - | 3864.1969 | 3.8-4.1 | 3774.51 |
| 5G2d | 0 | - | 8496.082 | 3.6-3.9 | 8224.76 |

Table S2

| Tukey's multiple comparisons test | Summary | Adjusted P Value |
|---|---|---|
| 5G0d (4 min) vs. 5G0d (5 min) | **** | <0.0001 |
| 5G0d (4 min) vs. 5G0.5d (1 min) | ns | 0.1721 |
| 5G0d (4 min) vs. 5G0.5d (2 min) | **** | <0.0001 |
| 5G0d (4 min) vs. 5G0.5d (3 min) | **** | <0.0001 |
| 5G0d (4 min) vs. 5G1d (30 sec) | ns | 0.0795 |
| 5G0d (4 min) vs. 5G1d (1 min) | **** | <0.0001 |
| 5G0d (4 min) vs. 5G1d (2 min) | **** | <0.0001 |
| 5G0d (4 min) vs. 5G2d (30 sec) | **** | <0.0001 |
| 5G0d (4 min) vs. 5G2d (1 min) | **** | <0.0001 |
| 5G0d (5 min) vs. 5G0.5d (1 min) | **** | <0.0001 |
| 5G0d (5 min) vs. 5G0.5d (2 min) | ns | 0.4014 |
| 5G0d (5 min) vs. 5G0.5d (3 min) | **** | <0.0001 |
| 5G0d (5 min) vs. 5G1d (30 sec) | **** | <0.0001 |
| 5G0d (5 min) vs. 5G1d (1 min) | ns | 0.0795 |
| 5G0d (5 min) vs. 5G1d (2 min) | **** | <0.0001 |
| 5G0d (5 min) vs. 5G2d (30 sec) | ns | 0.9095 |
| 5G0d (5 min) vs. 5G2d (1 min) | **** | <0.0001 |
| 5G0.5d (1 min) vs. 5G0.5d (2 min) | ns | 0.0666 |
| 5G0.5d (1 min) vs. 5G0.5d (3 min) | **** | <0.0001 |
| 5G0.5d (1 min) vs. 5G1d (30 sec) | ns | >0.9999 |
| 5G0.5d (1 min) vs. 5G1d (1 min) | ns | 0.3603 |
| 5G0.5d (1 min) vs. 5G1d (2 min) | **** | <0.0001 |
| 5G0.5d (1 min) vs. 5G2d (30 sec) | **** | <0.0001 |
| 5G0.5d (1 min) vs. 5G2d (1 min) | **** | <0.0001 |
| 5G0.5d (2 min) vs. 5G0.5d (3 min) | **** | <0.0001 |
| 5G0.5d (2 min) vs. 5G1d (30 sec) | ns | 0.1482 |
| 5G0.5d (2 min) vs. 5G1d (1 min) | ns | 0.9992 |
| 5G0.5d (2 min) vs. 5G1d (2 min) | **** | <0.0001 |
| 5G0.5d (2 min) vs. 5G2d (30 sec) | ** | 0.0087 |
| 5G0.5d (2 min) vs. 5G2d (1 min) | **** | <0.0001 |
| 5G0.5d (3 min) vs. 5G1d (30 sec) | **** | <0.0001 |

| | | |
|---|---|---|
| 5G0.5d (3 min) vs. 5G1d (1 min) | **** | <0.0001 |
| 5G0.5d (3 min) vs. 5G1d (2 min) | ns | 0.3791 |
| 5G0.5d (3 min) vs. 5G2d (30 sec) | **** | <0.0001 |
| 5G0.5d (3 min) vs. 5G2d (1 min) | **** | <0.0001 |
| 5G1d (30 sec) vs. 5G1d (1 min) | ns | 0.5708 |
| 5G1d (30 sec) vs. 5G1d (2 min) | **** | <0.0001 |
| 5G1d (30 sec) vs. 5G2d (30 sec) | **** | <0.0001 |
| 5G1d (30 sec) vs. 5G2d (1 min) | **** | <0.0001 |
| 5G1d (1 min) vs. 5G1d (2 min) | **** | <0.0001 |
| 5G1d (1 min) vs. 5G2d (30 sec) | *** | 0.0005 |
| 5G1d (1 min) vs. 5G2d (1 min) | **** | <0.0001 |
| 5G1d (2 min) vs. 5G2d (30 sec) | **** | <0.0001 |
| 5G1d (2 min) vs. 5G2d (1 min) | ns | 0.2561 |
| 5G2d (30 sec) vs. 5G2d (1 min) | **** | <0.0001 |
